# Lifting the ban on nuclear import activates Gdown1-mediated modulation of global transcription and facilitates adaptation to cellular stresses

**DOI:** 10.1101/2022.03.29.486178

**Authors:** Zhanwu Zhu, Jingjing Liu, Huan Feng, Yanning Zhang, Ruiqi Huang, Qiaochu Pan, Jing Nan, Ruidong Miao, Bo Cheng

## Abstract

Dynamic regulation of transcription is crucial for cellular response to various environmental or developmental cues. Gdown1 is a ubiquitously expressed, RNA polymerase II (Pol II) interacting protein, essential for embryonic development. It tightly binds Pol II *in vitro* and competitively blocks binding of TFIIF and other transcriptional regulatory factors, yet its cellular functions and regulatory circuits remain unclear. Here, we show that Gdown1 strictly localizes in the cytoplasm of mammalian somatic cells and exhibits potent resistance to the imposed driving force for nuclear localization. Combined with genetic and microscope-based approaches, two types of functionally coupled and evolutionally conserved localization regulatory motifs are identified, including the CRM1-dependent nucleus export signal (NES) and a novel Cytoplasm Anchoring Signal (CAS) which mediates nuclear pore retention. Mutagenesis of CAS alleviates the cytoplasmic retention activity thus unlocks its nucleocytoplasmic shuttling properties, and increased nuclear import of Gdown1 causes drastic reduction of Pol II levels and global transcription. Importantly, nuclear translocation of Gdown1 occurs in a stress-responsive manner and ablation of *GDOWN1* significantly weakens cellular tolerance. Collectively, our work uncovers the molecular basis of the localization of Gdown1 and highlights that its controlled nuclear translocation serves as a key strategy in modulating global transcription and stress-adaptation.

## INTRODUCTION

In eukaryotes, RNA Polymerase II (Pol II) catalyzes the RNA synthesis of all protein coding genes and a great number of non-coding genes in eukaryotic genomes(Haberle and Stark, 2018; Osman and Cramer, 2020). The core of transcription machinery is composed of the 12-subunit Pol II and a collection of dynamically bound and delicately coordinated factors, including general transcription factors (TFIID, TFIIA, TFIIB, TFIIF, TFIIE, TFIIH and TFIIS)(Cramer, 2019; Fischer et al., 2019), Pol II processivity-controlling factors (such as the writer, reader and eraser factors for modifying and recognizing the carboxyl terminal domain (CTD) of Rbp1(Hsin and Manley, 2012; Jeronimo et al., 2016; Sanso and Fisher, 2013; Yurko and Manley, 2018), the positive or negative elongation or termination factors)(Core and Adelman, 2019; Jonkers and Lis, 2015; Proudfoot, 2016; Zhou et al., 2012) and the co-transcriptional RNA processing and modifying factors (such as capping enzymes, splicing machinery, RNA modification enzymes, RNA cleavage factors) etc(Kachaev et al., 2020; Kilchert and Vasiljeva, 2013; Neugebauer, 2019; Noe Gonzalez et al., 2021; Schier and Taatjes, 2020; Sun et al., 2020). Many of these Pol II-binding and regulatory factors are not only essential for facilitating the production and processing of transcripts, but also play critical roles in dynamic integration of intracellular and extracellular information and adjusting Pol II’s target specificity and enzymatic activities in real time to maintain cell identity and homeostasis(Lynch et al., 2018; McNamara et al., 2016; Muniz et al., 2021; Schier and Taatjes, 2020).

Gdown1 was initially identified as a protein copurified with Pol II in calf thymus and porcine liver and designated as the 13^th^ subunit of Pol II due to its high-affinity interaction to Pol II (Hu et al., 2006). Data from *in vitro* transcription assays along with EMSA and structural analyses have demonstrated that Gdown1 strongly inhibits the binding and/or functions of a series of transcription regulatory factors, including the factors required for transcription initiation, such as TFIIF (Jishage et al., 2012), the factors involved in productive elongation RTF1/PAF1C (Ball et al., 2022), and the transcription termination factor such as TTF2 (Cheng et al., 2012).

Although the biochemical properties support Gdown1’s potential in regulating Pol II transcription, it is largely unknown how exactly its regulatory activities are executed under physiological circumstances. Knockout (KO) of *Gdown1* in flies and mice caused embryonic lethality, and moreover, the attempt of establishing a *Gdown1*-KO mouse ES cell line was failed, pointing out its essential roles during embryonic development (Jishage et al., 2020; Jishage and Roeder, 2020; Jishage et al., 2018). Interestingly, Gdown1 has been reported as a nucleocytoplasmic shuttling protein in flies. It colocalizes with Pol II in the nuclei at the transcriptionally silent syncytial blastoderm stage and moves to the cytoplasm at later blastoderm stage when global transcription is initiated, and similarly, Gdown1 is clearly retained in the nuclei of the transcriptionally silent pole cells (Jishage and Roeder, 2020). Although not being proved yet, these findings strongly suggest that controlling the nuclear import and export of Gdown1 is a key to make a switch of its transcription regulatory activities. At the beginning of embryonic development, the nuclear-localized Gdown1 may serve as a global transcription inhibitor, and once the embryo gets adequately prepared, exclusion of Gdown1 from the nucleus provides an effective way to promote zygotic genome activation (ZGA). Further studies are certainly required to explore the functional and regulatory mechanisms behind and find out whether these phenomena revealed in flies are similarly applied in higher animals or in other situations.

The piling up evidence suggest that Gdown1 also plays critical roles in somatic cells. It is expressed throughout the whole life cycle of flies (Jishage et al., 2018) and ubiquitously present across various types of mouse tissues (data not shown). Mice with Gdown1 specifically knocked out in liver were found to be viable and relatively normal, yet tended to trigger the quiescent hepatocytes to re-enter cell cycle in the absence of hepatic injury, highlighting its important role in maintaining the homeostasis of hepatocytes (Jishage et al., 2020). Further ChIP-Seq and RNA-Seq analyses revealed that Gdown1 bound with the elongating Pol II at many genes highlighting a collection of highly expressed genes in liver, while unexpectedly, Gdown1 showed a positive effect on transcription since the ablation of Gdown1 reduced Pol II occupancy and the transcription level of those genes (Jishage et al., 2020). Thus, it is necessary to clarify the underlying reasons behind the apparently opposite transcriptional regulatory effects of Gdown1 observed in somatic cells and in the defined *in vitro* transcription assays.

To further explore Gdown1’s functions in somatic cells, we started out by examining its subcellular localization in many cultured human cell lines and confirmed that it was predominantly localized in the cytoplasm. Based on the known functions, it’s reasonable to presume that GDOWN1 is a nucleocytoplasmic shuttling protein in somatic cells. However, our data demonstrated that GDOWN1 was subjected to very tight restriction for its nuclear import under the regular cell culture conditions. Based on these findings, we established various mutagenesis-based screening assays and identified multiple intrinsic localization regulatory signals and their working mechanisms. In addition, manipulation of GDOWN1’s nuclear translocation caused significant reduction of both Pol II protein level and the global transcription and its massive and constant nuclear accumulation caused severe growth inhibition and even triggered cell death. In addition, we provided evidence that the nuclear import of GDOWN1 was naturally induced upon certain cellular stresses and its genetic ablation was associated with reduced cell viability in stress response. Overall, our data revealed a novel function of Gdown1 in facilitating cellular adaptation to stresses via modulation of transcription homeostasis, and the execution of this protective strategy was associated with its controlled nuclear import.

## RESULTS

### Gdown1 is primarily a cytoplasm-localized protein in mammalian somatic cells

We started out to detect the subcellular localization of GDOWN1 in cultured human cell lines by ectopically expressing GDOWN1 fused with a fluorescent tag at its N- or C-terminus or simply with a Flag tag. Consistent to the previous observation in adult flies (Jishage et al., 2018) and the most recent report in human somatic cells (Ball et al., 2022), the localization signals of GDOWN1 were exclusively present in the cytoplasm of HeLa cells, regardless of the position or size of the fused tags (Fig. 1A). To explore GDOWN1’s functions in the nucleus, two nucleus localization signals (NLS) were fused to GDOWN1 at each end, which were known to be efficiently driving the 160 KDa SpCas9 protein into the nucleus in a commonly used CRISPR-vector pX459 (Ran et al., 2013). Unexpectedly, addition of two NLS motifs did not affect GDOWN1’s subcellular localization at all (Fig.1A). We then detected the nucleocytoplasmic distribution of endogenous GDOWN1 in fractionated cell lysates from various human and mouse cell lines by Western blotting using KO-verified Gdown1 antibodies (Fig. S1A). The results clearly indicated that endogenous Gdown1 was predominantly located in the cytoplasmic fractions in all the five human and mouse somatic cell lines tested and only a small fraction of Gdown1 was seen in the nuclear extract of mouse embryonic stem cells, E14TG2a (Figs. 1B, S1B). These data indicate that GDOWN1 is a strictly cytoplasm-localized protein in various human and mouse somatic cells.

**Figure 1.**
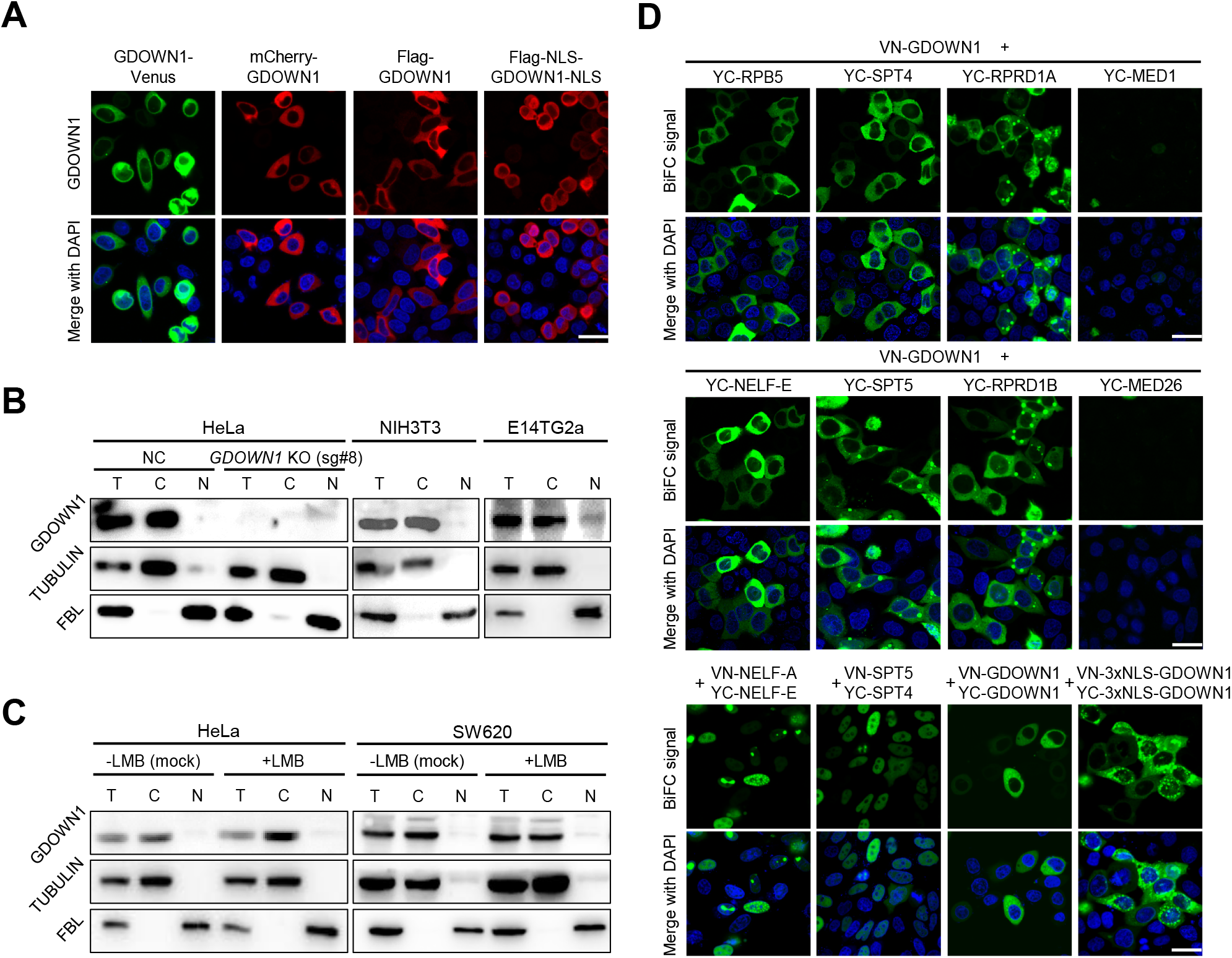
Detection of the subcellular localization of GDOWN1 or BiFC signal between GDOWN1 and some transcription related factors. **A.** The ectopically expressed human GDOWN1 in HeLa cells was stringently localized in the cytoplasm. Human GDOWN1 proteins fused with indicated tags, including a fluorescent tag at either terminus, a Flag tag alone or together with two NLS motifs, were ectopically expressed in HeLa cells and the subcellular localization was detected by directly monitoring the fluorescent signal or by immunofluorescence assays (IF) using an anti-FLAG antibody. **B.** The endogenous human or mouse GDOWN1 was stringently located in the cytoplasm. Each indicated cell line was fractionated to separate cytosol from nuclei and the cytoplasmic fraction (C), the nuclear fraction (N) and the whole cell lysate (T, total) were further detected by Western blot analyses (WB). α-TUBULIN and FBL (a nucleolus protein) were used as markers of the cytoplasmic and nuclear fractions, respectively. **C.** GDOWN1 remained in the cytoplasm upon LMB treatment. The indicated cell lines were subjected to either mock or LMB treatment (detailed below) before further fractionation and WB analyses. **D.** BiFC analyses of the protein-protein interactions between GDOWN1 and its potential binding partners. Proteins of interest were cloned and fused with either VN (the N-terminus of Venus) or YC (the C-terminus of YFP) and each indicated pair of plasmids were co-transfected into HeLa cells and the confocal microscopy images were acquired 24 hours post transfection. The LMB treatment was carried out at 20 nM final concentration for 6 hours and the mock treatment was done with an equal volume of ethanol in parallel. Nuclear DNA was stained by Hoechst 33342 and all the scale bars represent 30 μm. Without further labeled with details, the Gdown1 antibody used in WB assays were generated from rabbit.

It was known that mammalian Gdown1 interacted to Pol II very potently (Hu et al., 2006; Jishage et al., 2018) and data from *in vitro* transcription assays indicated that it had a Mediator-reversible inhibitory effect on Pol II transcription initiation (Cevher, 2021; Hu et al., 2006; Jishage et al., 2012) and may facilitate stabilizing the paused elongation complex (Cheng et al., 2012). Combining these potential nuclear functions and our observation of GDOWN1’s cytoplasmic localization, it is reasonable to hypothesize that maybe GDOWN1 is a nucleocytoplasmic shuttling protein that functions in the nucleus in a transient manner. Most of the nucleocytoplasmic shuttling proteins contain a nuclear export signal (NES) and the classical NES is known as a hydrophobic leucine-rich motif recognized by the ubiquitous transport receptor chromosome maintenance protein 1, CRM1 (also namely exportin 1) (la Cour et al., 2004; Xu et al., 2010). To test the possibility of GDOWN1 being a CRM1 cargo, HeLa and SW620 cells were treated with leptomycin B (LMB), a known CRM1 inhibitor for efficiently blocking CRM1-NES interaction (Kudo et al., 1999; Kudo et al., 1998). Western blotting using two KO-verified Gdown1 antibodies unambiguously demonstrated that GDOWN1 didn’t accumulate in the nucleus upon LMB treatment (Figs. 1C and S1C). The resistance to LMB treatment implies that either GDOWN1 does not have an NES, or this treatment by itself is insufficient to achieve nuclear accumulation of GDOWN1.

On the other hand, we employed BiFC assays to detect the interactions between GDOWN1 and its potential nuclear binding partners in live cells. An efficient interaction between the proteins of interest drives the formation of the fluorescence complementation, achieved via the covalent interactions between the two truncated parts of a fluorescent protein. Therefore, BiFC signals are irreversible once generated, making this assay beneficial of capturing transient protein-protein interactions. A series of transcription-related proteins were tested in HeLa cells, including Pol II subunit (RPB5) and the RPB1-CTD binding factors (RPRD1A, RPRD1B), the Mediator components (MED1, MED26), and transcription elongation factors (SPT4 and SPT5 in DSIF complex, NELF-E in NELF complex). As shown in Figure 1D, GDOWN1 interacted to all the above factors tested except for the two Mediator components, well supporting its known characters as a Pol II-associating factor and the potential functions involved in transcriptional regulation. However, all these BiFC signals were shown in the cytoplasm, yet in parallel tests the interaction signals between NEFL-E•NEFL-A, SPT4•SPT5 pairs were both exclusively present in the nucleus as expected. Meanwhile our BiFC assays detected the self-interaction of GDOWN1 in the cytoplasm, suggesting that GDOWN1 may form homodimers or oligomers in cells. It was reported that transcription regulator RYBP contained three potent and functionally independent NLSs (Tan et al., 2017) and when attached to GDOWN1, the BiFC signal of the 3xNLS-GDOWN1 dimers mainly remained in the cytoplasm (Fig. 1D). These results support GDOWN1’s potential functions in transcriptional regulation while the stringent cytoplasmic localization of the BiFC signals indicates that GDOWN1 is restricted from entering the nucleus under normal cell culture conditions. Thus, our data further confirm that the nuclear entry of GDOWN1 is subjected to tight regulation and suggest that alleviation of this restriction is a prerequisite for permitting GDOWN1’s nuclear functions in transcriptional regulation.

### Gdown1’s cytoplasm-localization is determined by two distinct types of localization regulatory signals

Next, we constructed a series of GDOWN1 mutants to screen for localization regulatory signals by monitoring the changes of the subcellular localization of themselves or together with other proteins. LMB treatment was applied to further analyze the possibility of containing NES. Consistent to the above cell fractionation results, both the ectopically expressed full length GDOWN1, and the BiFC signal of GDOWN1 and NELF-E remained in the cytoplasm in the presence of LMB (Figs. 2A-B, S2A). Then GDOWN1 was truncated into three parts at its structurally flexible regions (N-terminus, mutant #1, namely *m1*; middle part, *m2*; C-terminus, *m3*), fused with Flag-VN in BiFC vector or fused with Venus to monitor their dynamic localization in the absence or presence of LMB treatment (Figs. 2A-B). GDOWN1-*m1* was mainly located in the nucleus (Figs. 2B, left panel; S2A) and the *m1*•NELF-E BiFC signal was completely nucleus localized (Fig. 2B, right panel). The other two counterparts, *m2* and *m3* remained their own subcellular localization and interacted to NELF-E in the cytoplasm. Interestingly, both *m2* alone and *m2*•NELF-E signals were translocated into the nucleus in response to LMB, while either *m3* alone or the *m3*•NELF-E signal did not respond to LMB at all (Figs. 2B and S2A). The consistent results obtained from direct or indirect detection clearly indicate that the middle part of GDOWN1 contains an NES motif. Given that the translocation of GDOWN1 into nucleus may not be an autonomous and efficient process, we reasoned that monitoring the nuclear accumulation of BiFC signals between GDOWN1 and its nuclear binding partners (such as NELF-E) had the advantage for better mining and demonstrating the nucleus translocation potential of GDOWN1. Thus, the above BiFC system was further employed for screening the putative localization regulatory motif(s). When *m1* and *m2* parts were combined, the resultant fragment, *m4*, was able to respond to LMB as well as *m2* alone. The conserved sequence of a classical NES for CRM1 recognition was known as Ψ-(x)_1-3_-Ψ-(x)_1-3_-Ψ-(x)_1-3_-Ψ (Ψ stands for L, I, V, M, or F, x can be any amino acid) (la Cour et al., 2004; Xu et al., 2012), we tested a series of GDOWN1 truncation mutants to search for the functional NES within *m2* region (Fig. S2B), and further confirmed that a putative NES motif located between amino acids 191-201 was responsible for LMB response. Mutation of the four hydrophobic amino acids within this region completely abolished the NES activity (Fig.2A-B, *m4\**). By co-immunoprecipitation and BiFC assays, we confirmed that GDOWN1 interacted with CRM1/RAN, the core components for the protein nuclear export machinery (Fig. 2C). These results prove that GDOWN1 indeed contains a classical CRM1-dependent NES motif and meanwhile suggest that the C-terminus of GDOWN1 contains a regulatory motif responsible for the observed resistant activity of full-length GDOWN1 to LMB treatment.

**Figure 2.**
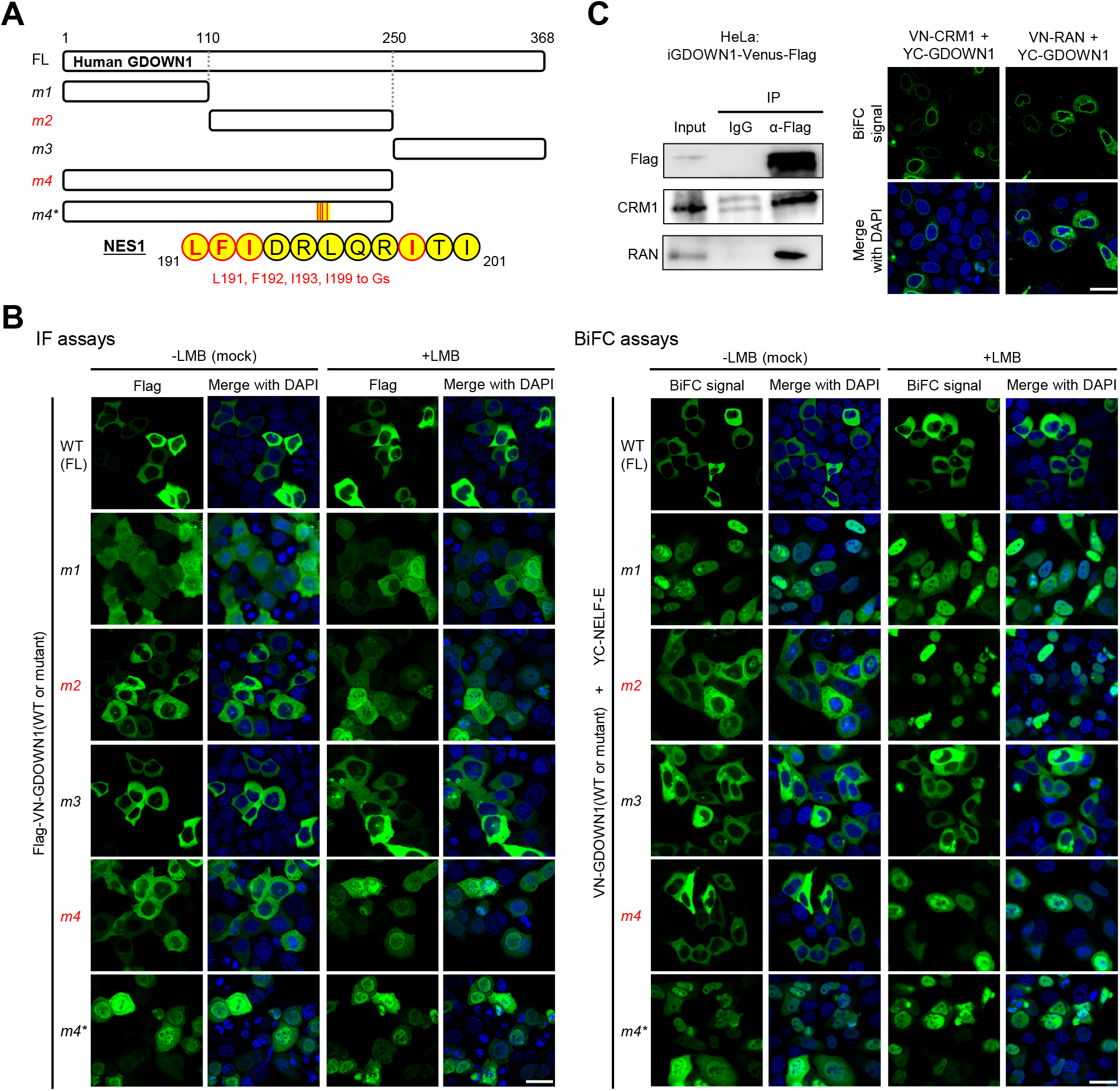
Identification of Nuclear Export Signal (NES) motifs in GDOWN1. **A.** A diagram of human GDOWN1 and its mutants used in the IF or BiFC-based motif screening analyses. The mutants whose names are marked in red are the ones translocated into the nucleus in response to LMB treatment. The position and sequences of the identified NES motifs are shown in yellow circles and the core amino acids selected for mutagenesis are highlighted in red. **B.** Identification of the NES motifs in GDOWN1 via IF or BiFC-based screening analyses. Left panel: HeLa cells were transiently transfected with plasmid carrying Flag-WT or mutant GDOWN1 as indicated, and further subjected to either mock or LMB treatment, the subcellular localization was detected by IF using a Flag antibody; Right panel: HeLa cells were transiently transfected with two BiFC plasmids, YC-NELF-E and VN-WT or mutant GDOWN1 as indicated (VN—the N-terminus of Venus; YC—the C-terminus of YFP), and further subjected to either mock or LMB treatment before signal detection by a confocal microscope. **C.** Detection of the interaction between GDOWN1 and CRM1 or RAN by IP-WB or BiFC assays. Left panel: HeLa cells stably expressed GDOWN1-Venus-Flag were employed for IP experiment using a Flag antibody or IgG and further detected by WB with indicated antibodies; Right panel: BiFC analyses of GDOWN1•CRM1/RAN interactions. HeLa cells were transfected with YC-GDOWN1 and VN-CRM1 or RAN. The LMB treatment was carried out at 20 nM final concentration for 6 hours and the mock treatment was done with an equal volume of ethanol in parallel. Nuclear DNA was stained by Hoechst 33342 and all the scale bars represent 30 μm.

When *m1* and *m3* parts were combined to generate *m5*, it was not subjected to LMB-dependent nuclear accumulation but became LMB-responsive when the very end of the C-terminus was chopped off, which led to the identification of the second NES (Figs.3A-B and S3A, *m5*, *m6*). Further mutant screening identified the second NES located between amino acids 332-340 (Figs. S3B-C, *m12, m13* and Figs. 3A-B and S3A, *m6\**). Taken together, we confirm that GDOWN1 is a CRM1 cargo containing two classical CRM1-responsive NES.

**Figure 3.**
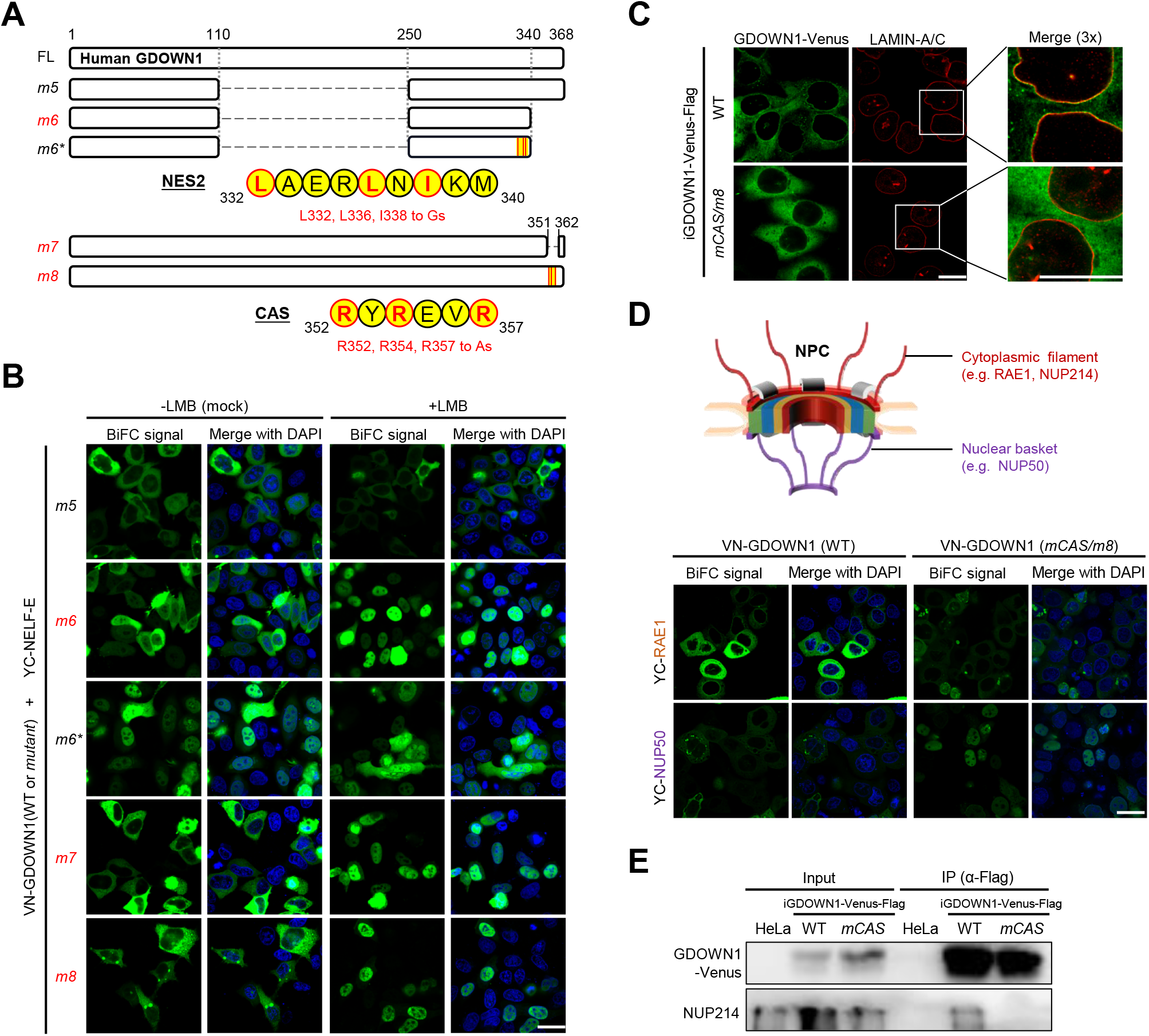
Identification and mechanism analysis of the Cytoplasm Anchoring Signal (CAS) motif in GDOWN1. **A.** A diagram of human GDOWN1 and its mutants used in the BiFC-based motif screening analyses. The mutants whose names are marked in red are the ones translocated into the nucleus in response to LMB treatment. The position and sequences of the identified NES or CAS motif are shown in yellow circles and the core amino acids selected for mutagenesis are highlighted in red. **B.** Identification of the second NES and the CAS motifs in GDOWN1 via BiFC-based screening analyses. The experiments were carried out in the same way as described in Figure 2B. **C.** The enrichment of GDOWN1 at the nuclear pore region was regulated by the CAS motif. HeLa cells stably expressing the wild type GDOWN1 (WT-Venus) or the CAS mutant (*mCAS*-Venus) were used for detection. The nuclear membrane was approximately represented via IF using an antibody against the nuclear lamina (α-LAMIN-A/C). Confocal Images were collected and further zoomed in for 3 folds to show more details of the nuclear membranes. **D.** BiFC analyses of the interactions between GDOWN1 and some subunits of NPC (nuclear pore complex) in HeLa cells. Upper panel: a simplified diagram of an NPC; lower panel: BiFC results between GDOWN1 (or its CAS mutant) and the indicated NPC components. **E.** Detection of the interaction between GDOWN1 and NUP214 by IP-WB. Parental Hela cells or HeLa cells stably expressed GDOWN1(WT or *mCAS*)-Venus-Flag were employed in IP experiment using a Flag antibody and further detected by WB with indicated antibodies. The LMB treatment was carried out as previously described. Nuclear DNA was stained by Hoechst 33342. All scale bars in this figure represented 30 μm, except for the ones labeled in C represented 15 μm.

The distinct responsiveness of the *m5* and *m6* parts of GDOWN1 to LMB treatment clearly indicated that the C-terminus of GDOWN1 contained another CRM1-independent, cytoplasmic localization regulatory signal. The key amino acids responsible were then examined in the BiFC reporter system by screening the C-terminal truncation or deletion mutants (Figs. S3B-C, *m14-16*). It turned out that deletion of amino acids 352-361 abolished this cytoplasm localization regulatory activity and switched GDOWN1 from LMB-irresponsive to LMB-responsive manner (Figs. 3A-B and S3A, *m7*). After testing a series of combinations of point mutations, we found mutations of the three arginines (R352, R354, R357) were efficient to abolish the above cytoplasm localization activity of GDOWN1 in the presence of LMB (Figs. S3B-C, *m17*, *m18*; Figs. 3A-B and S3A, *m8*). Due to its potent cytoplasmic retention activity, we named this region (352-357 aa) Cytoplasm Anchoring Signal, CAS.

To further elucidate the working mechanism of the CAS motif, we generated stable HeLa cell lines that inducibly expressed either the wild type GDOWN1 (WT-Venus) or its CAS mutant (*mCAS*-Venus) (Fig. S4A, top). In these stable cell lines, the dynamic localizations of GDOWN1 in the presence or absence of LMB were monitored and the results were consistent to those obtained from the above transient transfection assays (Fig. S4A, bottom). Interestingly, the cytoplasmic localization of the wild type GDOWN1 and CAS mutant was obviously different in the high-resolution confocal microscopy images. The wild type GDOWN1 accumulated around the nuclear membrane, as if these molecules attempted to burst through this last defense line for their nucleus entry, while the CAS mutant lost this “ring-form accumulation” surrounding the nuclear membrane, and became widely scattered all over the cytoplasm (Fig. 3C). We hypothesized that the Venus signal enriched around the nuclear membrane was an indicator of GDOWN1 associated with the Nuclear Pore Complex (NPC). Due to the complicated composition of NPC, we detected the interaction of GDOWN1 to representative NPC components via BiFC assays. RAE1 and NUP50 are two NPC components typically assembled within the cytoplasmic filaments and the nuclear baskets, respectively. BiFC results indicated that wild type GDOWN1 strongly interacted to RAE1 at nuclear membrane and in the cytoplasm while this interaction was drastically weakened in the CAS mutant (Fig. 3D), suggesting that the CAS motif was involved in GDOWN1–NPC interaction. More interestingly, the BiFC signal of wild type GDOWN1 and NUP50 was very weak and randomly distributed throughout the cytoplasm while the CAS mutant specifically translocated this binding signal into the nucleus, especially at the inner face of the nuclear membrane where NUP50 naturally located (Figs. 3D, and S4B). IP results also demonstrated that the wild type GDOWN1 interacted with the cytoplasmic NPC component, NUP214, while the CAS mutant lost this interaction (Fig. 3E). Overall, these results demonstrated that wild type GDOWN1 specifically interacted to the cytoplasmic NPC components, while the CAS mutant reduced this binding affinity and simultaneously enhanced the interaction of GDOWN1 to the nuclear NPC components. Due to the irreversible nature of BiFC signal, the nuclear signal of *mCAS*-GDOWN1•NUP50 interaction was a clear indication of successful capture of this GDOWN1 mutant in the nucleus, while its wild type counterpart was restricted in the cytoplasm. The above data highlight the crucial role of the CAS motif on locking GDOWN1 in the cytoplasm, presumably through anchoring of GDOWN1 to the cytoplasmic components of NPC, and imply that any cellular strategy of preventing CAS from functioning will switch GDOWN1 from a stringent cytoplasm-localized protein into a nucleocytoplasmic-shuttling protein.

### The NES and CAS motifs in Gdown1 are functionally interconnected and both conserved during evolution

Based on the structural prediction of GDOWN1, its CAS motif is located within the disordered region near the carboxyl-terminus, which makes it difficult to obtain reliable structural information to predict the potential CAS-NES interaction (Fig. S4C). Indeed, a previous report carrying out chemical crosslinking with mass spectrometry readout (CX-MS) to analyze Gdown1-Pol II interaction did not provide any information about its C-terminal CAS region (Jishage et al., 2018). To clarify the functional relationship between CAS and NES motifs, we transiently expressed GDOWN1-Venus or its localization motif-mutants that carried combinations of the key amino acid mutations identified above, and tested their subcellular localization and LMB responsiveness (Fig. 4A). When both NES2 and CAS were mutated to allow NES1 alone to function, the resultant GDOWN1 mutant performed as a classical CRM1-cargo and on the other hand, the NES1 mutant maintained the same cytoplasmic localization and LMB resistance activity as the wild type GDOWN1 (Figs. 4A-B, a-c). Thus, NES1 was a functional NES motif working independently but redundantly to NES2. When NES2 alone was remained, although its cytoplasm localization became less stringent, this mutant responded to LMB treatment very well so that it was a functional NES as well (Figs. 4A-B, d). The NES2 mutant remained its cytoplasm localization regularly, but did not resist to LMB treatment as well as the wild type, suggesting that the cytoplasm localization activity of CAS might be partially interfered in this NES2 mutant (Figs. 4A-B, e). Double mutations in both NES motifs made GDOWN1 distributed in both the cytoplasm and the nucleus, and did not respond much to LMB, which proved that the entire GDOWN1 contained two NES motifs and again mutations of NES2 partially abolished CAS activity (Figs. 4A-B, f). Comparing to the wild type, the CAS mutant responded well to the LMB treatment, strengthening the point that the CAS motif anchor GDOWN1 in the cytoplasm in a CRM1-independent manner, while it had to execute this activity in concert with the NES2 region (Figs. 4A-B, g). The above data from the intrinsic motif analyses demonstrate that each one of the two NES motifs of GDOWN1 acts as an independent CRM1-regulated element and the function of CAS motif depends partially on the structural support from NES2, but not rely on its CRM1 binding activity. Taken together, GDOWN1 is identified as a nucleocytoplasmic shuttling protein subjected to both CRM1-dependent and CRM-independent regulation and the two layers of regulation are interconnected.

**Figure 4.**
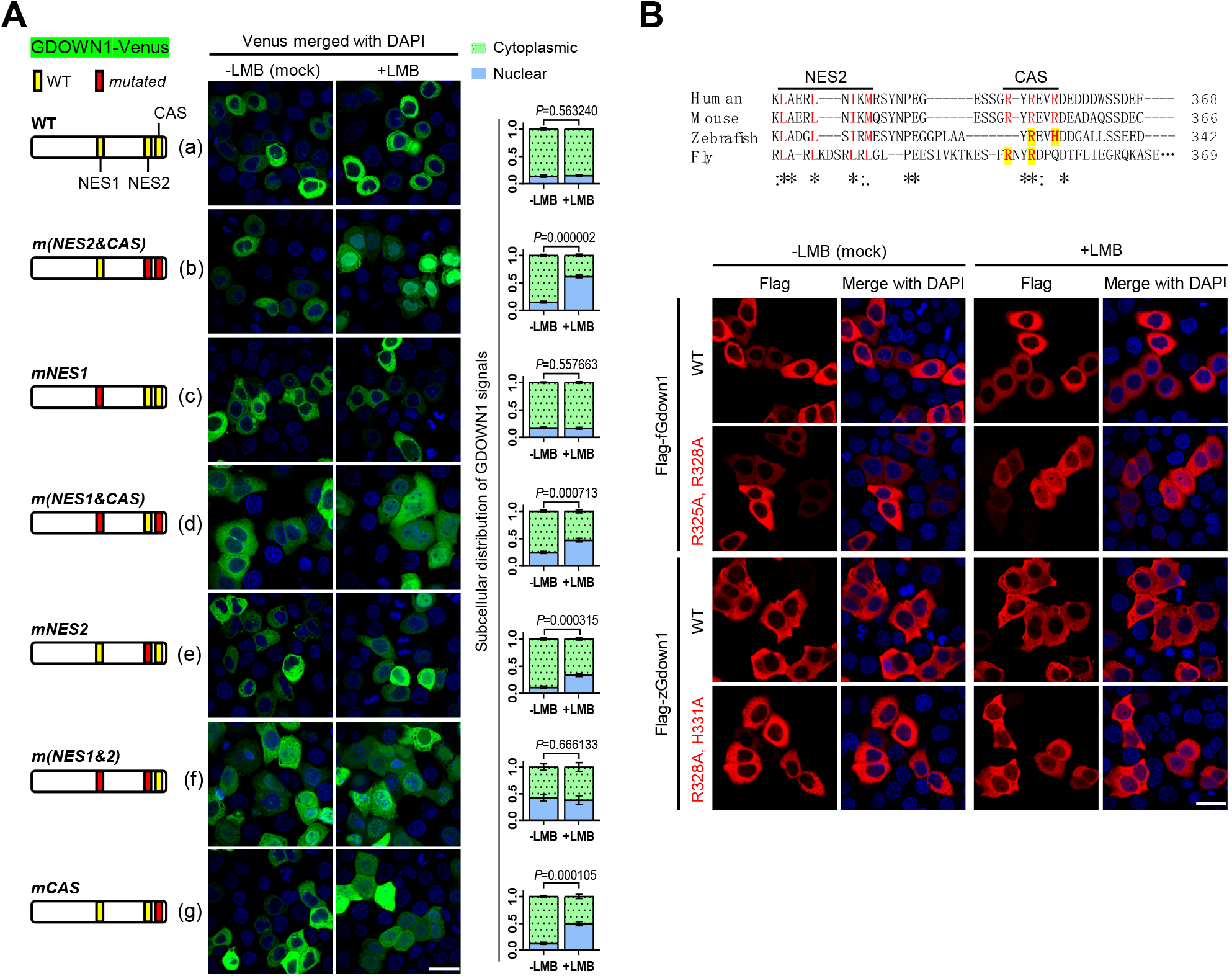
The working mechanisms and conservation of the binary localization regulatory apparatus in Gdown1. **A.** Dissection of the functional independence and interplay among CAS and NES motifs. Wild type GDOWN1 or the indicated CAS or NES mutants carrying point mutations were fused with Venus and ectopically expressed in HeLa cells. The cells were subjected to mock or LMB treatment the same as described in Figure 1. The schematic diagram of each mutant is shown on the left side of the corresponding representative confocal microscopy images. The nucleocytoplasmic distribution of the fluorescent signals was quantified using ImageJ and shown on the right. For statistics analyses, the calculated values were further processed to obtain the *P* values via t-test using the built-in tools in Graphpad Prism8. **B.** The function of the NES and CAS motifs was very conservative from zebrafish and drosophila to mammals. Upper panel: sequence alignment of the putative NES2-CAS regions of Gdown1 proteins from the indicated species (Homo sapiens, NP_056347.1, Mus musculus, NP_848717.1, Danio rerio, NP_001333109.1, Drosophila melanogaster, NP_650794.1). “**\***”—identical in all species analyzed; “:”—highly conserved; “.”— moderately conserved. Lower panel: the dynamic subcellular localization of the wild type or CAS mutants of zebrafish (zGdown1) and fly (fGdown1) was detected by IF experiments. The plasmids expressing indicated proteins were transfected into HeLa cells and the LMB treatment was carried out as previously described. scale bars—30 μm.

Since the nucleocytoplasmic-shuttling effect of Gdown1 was reported in drosophila, we evaluated the conservation of its localization regulatory mechanisms across species. The sequences corresponding to the three localization regulatory motifs in Gdown1 from various representative species (fly, zebrafish, mouse and human) were compared via Clustal Omega analyses. The NES motifs are modestly conserved across these species with the key hydrophobic amino acids roughly present in fly and zebrafish Gdown1 proteins (Figs. S4D and 4B). In terms of the CAS motifs, there is no difference between mouse and human, while there is only one or two key arginines remained present in the putative CAS motifs of zebrafish and fly Gdown1 proteins, respectively. When ectopically expressed in HeLa cells, fly and zebrafish Gdown1 proteins also located stringently in the cytoplasm and resisted to LMB treatment as same as their human counterpart (Fig. 4B). When the conserved amino acids in putative CAS motifs of fly and zebrafish Gdown1 proteins were mutated, these mutants became partially nucleus localized upon LMB treatment (Fig. 4B), indicating that fly and zebrafish Gdown1 also contained functional NES and CAS motifs. In addition, results from BiFC analyses demonstrated that fly and zebrafish Gdown1 proteins were able to interact to human NELF-E in the cytoplasm, indicating that these orthologs in lower animals were structurally conservative to human GDOWN1 (Fig. S4E). Different from the mammalian counterpart, the BiFC signals between fly or zebrafish Gdown1 and NELF-E were partially translocated into nucleus in the presence of LMB, and when CAS regions were mutated, these BiFC signals were completely present in the nucleus, suggesting that the regulatory effect of CAS in fly and zebrafish Gdown1 was not as potent as in human (Fig. S4E). The above results demonstrate that both the CRM1-dependent and CRM1-independent regulatory mechanisms of Gdown1 are well conserved across from flies to human, while during evolution, the cytoplasm anchoring effect of CAS motif seems to be gradually enhanced to strengthen the regulation of Gdown1’s subcellular localization.

### Nuclear-localized GDOWN1 modulates total Pol II level and the global transcription and its massive accumulation inhibits cell growth

The great effort devoted by the cells to prevent Gdown1 from entering the nucleus strongly implies that it is essential to stringently control the nuclear activities of Gdown1. To help explore the outcome of Gdown1’s nuclear accumulation in somatic cells, we set up to generate a nucleus-localized, full-length human GDOWN1 mutant by mutating all the ten key amino acids identified in the three motifs of NES and CAS (highlighted in red in Figures 2A and 3A, simply named *10M* mutant). The wild type or *10M* mutant GDOWN1 was fused with Venus and cloned into a commercial pTripZ vector to achieve doxycycline (Dox)-inducible expression and stable HeLa cell lines were generated. Figure 5A is a diagram of the experimental procedures. As demonstrated in Figure 5B-(i), the *10M* mutant was evenly distributed in cells and further addition of an NLS motif switched GDOWN1 into a complete nucleus localized protein (NLS-*10M*). These stable cell lines were generated by collecting the pool of cells survived from puromycin selection, which turned out to be heterogenous population that contained both Venus^+^ and Venus^-^ cells upon Dox induction. The benefit of using such heterogenous cell pools instead of the single clones hereby was that the co-cultured Venus^-^ cells (expressing none or very low level of GDOWN1-Venus) served as the internal negative controls for comparing cellular activities to the GDOWN1-highly expressed, Venus^+^ cells (Fig. 5B-i). When Dox was continuously supplemented in the culture medium, the fluorescence intensity in Venus^+^ cells and their ratio to the whole population reached nearly maximum around day 3 and remained stable hereafter in the two cells lines expressing the wild type GDOWN1 (Fig. 5B-ii). However, these values were significantly reduced in the two cell lines expressing the nucleus-localized *10M* mutants, especially expressing NLS-*10M*-Venus resulted in almost complete loss of the Venus signal on day 9 after the initial Dox addition, indicating that the accumulation of GDOWN1 in the nucleus was unfavorable for the cell growth (Fig. 5B-ii). To dissect the underlined reasons for signal loss, we comprehensively evaluated the growth status of the cells after inducing expression of either cytoplasm- or nucleus-localized GDOWN1 (withdrawal of Dox on day 3). The cells on day 0 (Dox addition), day 3 (Dox withdrawal) and day 9 were analyzed by FACS. The results demonstrated that cells expressing cytoplasm-localized GDOWN1 only showed basal level of cell death (indicated by the DAPI^+^ subgroup), while the cells expressing nuclear-localized GDOWN1-*10M* showed drastic reduction of expression (indicated by the decreased FITC values) and simultaneously those Venus^+^ cells mainly contributed to the significant increased death rate at later time point (Fig. 5B-iii). In addition, the results from the cell counting and the real-time cell analysis assays (RTCA) both demonstrated that the cells expressing *10M* mutant had severe defects on their growth and proliferation (Figs. 5B-iv, S5). The above data indicate that massive accumulation of GDOWN1 in the nucleus inhibits cell growth and the continuous accumulation eventually causes cell death.

**Figure 5.**
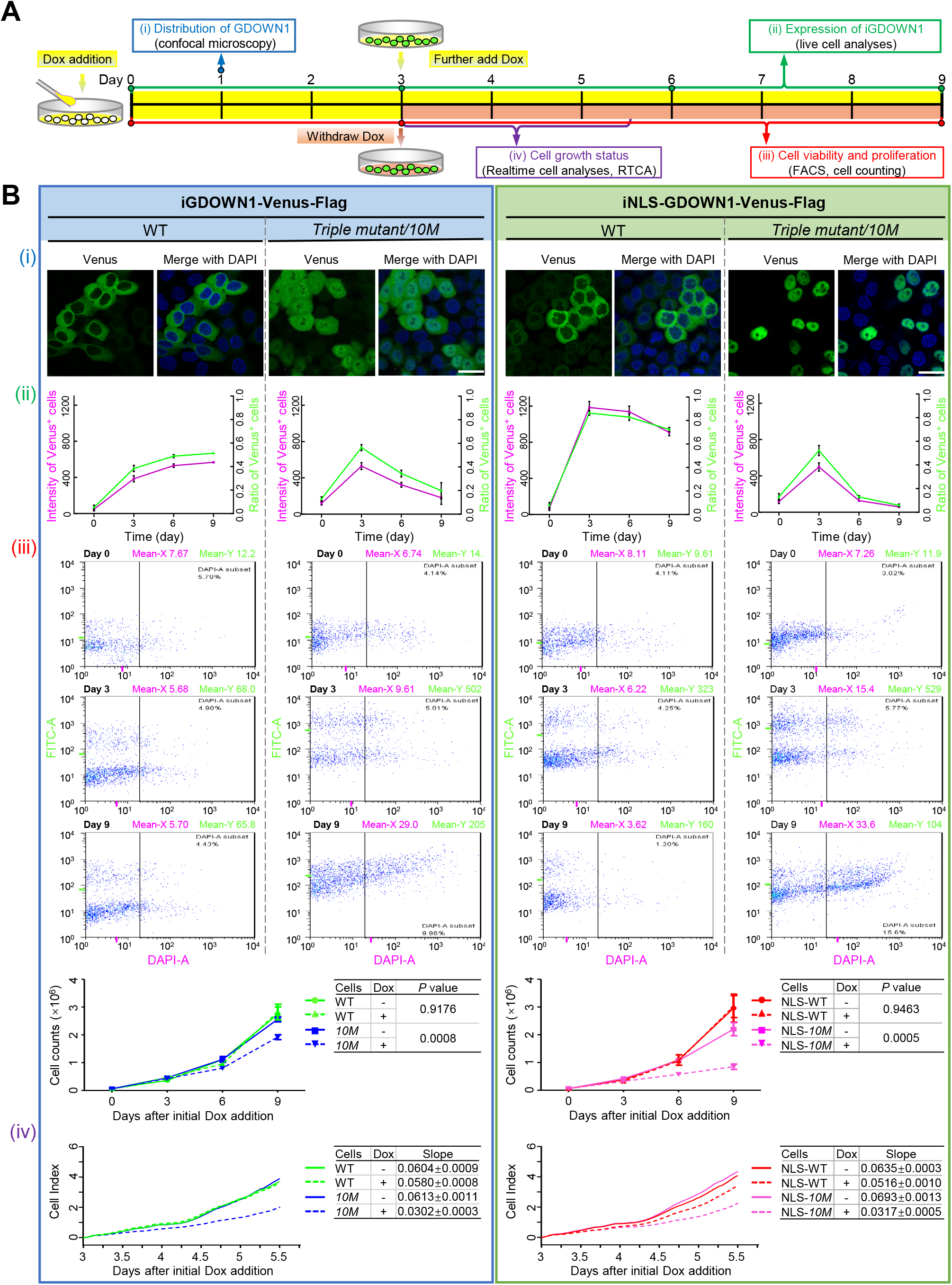
Massive accumulation of GDOWN1 in the nucleus slows down cell growth and may even trigger cell death. HeLa cells stably and inducibly expressing GDOWN1- or NLS_sv40_-GDOWN1-Venus-Flag, wild type or the triple mutant (*10M*) were used for detection, “i” stands for inducible and doxycycline was used as the inducer. **A.** The experimental scheme of the comprehensive analyses of the GDOWN1 expressing cell lines. **B. (i)** Confocal images demonstrating the subcellular localization of the indicated cells upon doxycycline induction for 1 day. Nuclear DNA was stained by Hoechst 33342. scale bars—30 μm. **(ii)** The changes of the fluorescence intensity and the ratio of Venus^+^ cells upon induction of GDOWN1 expression. Images were acquired by Cytation 5 and data were further analyzed by Gene 5. **(iii)** Monitor cell death and the changes of the fluorescence intensity via flow cytometry. Cells were induced by doxycycline for 3 days to reach maximum expression and continuously cultured for 6 days in the absence of doxycycline. Then, the cells were subjected to a quick DAPI staining, followed by flow cytometry analyses. The mean values of FITC signal (bright green, indicating the expression levels of GDOWN1-Venus proteins) and of the DAPI signals (pink) are labeled on each graph. The gating parameter for DAPI^+^ dead cells was set based on the readings of a control sample containing known ratio of live and dead cells and the portion of dead cells in each sample was shown. Meanwhile, cells were counted at 0, 3, 6 and 9 days and the growth curves are shown at bottom. **(iv)** Cell growth status monitored by a live cell analyzer. Cells cultured with doxycycline for 3 days were replated in the same cell number in a gold-coated 16-well plate for RTCA and further cultured in the presence or absence of doxycycline for 2.5 days. The real time cell index parameter was recorded and plotted by RTCA. The doxycycline was applied at a final concentration of 0.25 μg/mL.

It was known from *in vitro* transcription assays that Gdown1 negatively regulated Pol II transcription via competing TFIIF from binding, therefore we reasoned the cell death effects seen here might be resulted from GDOWN1-mediated transcriptional changes. EU incorporation assays were carried out using the above four cell lines expressing WT or *10M* GDOWN1 and the EU signals were pseudo-colorized based on the acquired intensity, which correlated to the overall transcription levels in each cell. It turned out that the expression of WT GDOWN1 did not cause obvious changes of transcription while in the cell lines expressing nuclear GDOWN1, the EU incorporation in the Venus^+^ cells significantly decreased comparing to the Venus^-^ cells, indicating that GDOWN1’s abundance in the nucleus negatively correlated with the overall extent of cellular transcription (Figs. 6A, and S6A). Next, we monitored the changes of Pol II in these cells by immunofluorescence assays and Pol II signals were detected using the pan antibody targeting to the total level of the largest subunit of Pol II, RPB1, or via the antibodies specifically recognizing its CTD-phosphorylated form at either Ser5 positions (S5P, detecting transcriptionally initiated Pol II) or Ser2 positions (S2P, detecting Pol II engaged in productive elongation). Comparing to the parental HeLa cells or stable cell lines expressing WT GDOWN1, the signals of the total and the phosphorylated forms of Pol II were all dramatically reduced in cell lines expressing nucleus-localized GDOWN1 (Figs. 6B, and S6B). Results from the cell fractionation and WB assays further indicated that the total protein levels of Pol II reduced upon nuclear accumulation of GDOWN1 while the nucleocytoplasmic ratio of Pol II seemed not affected. Taken together, these results demonstrate that massive nuclear translocation of GDOWN1 results in reduction of Pol II and global transcriptional shut-down.

**Figure 6.**
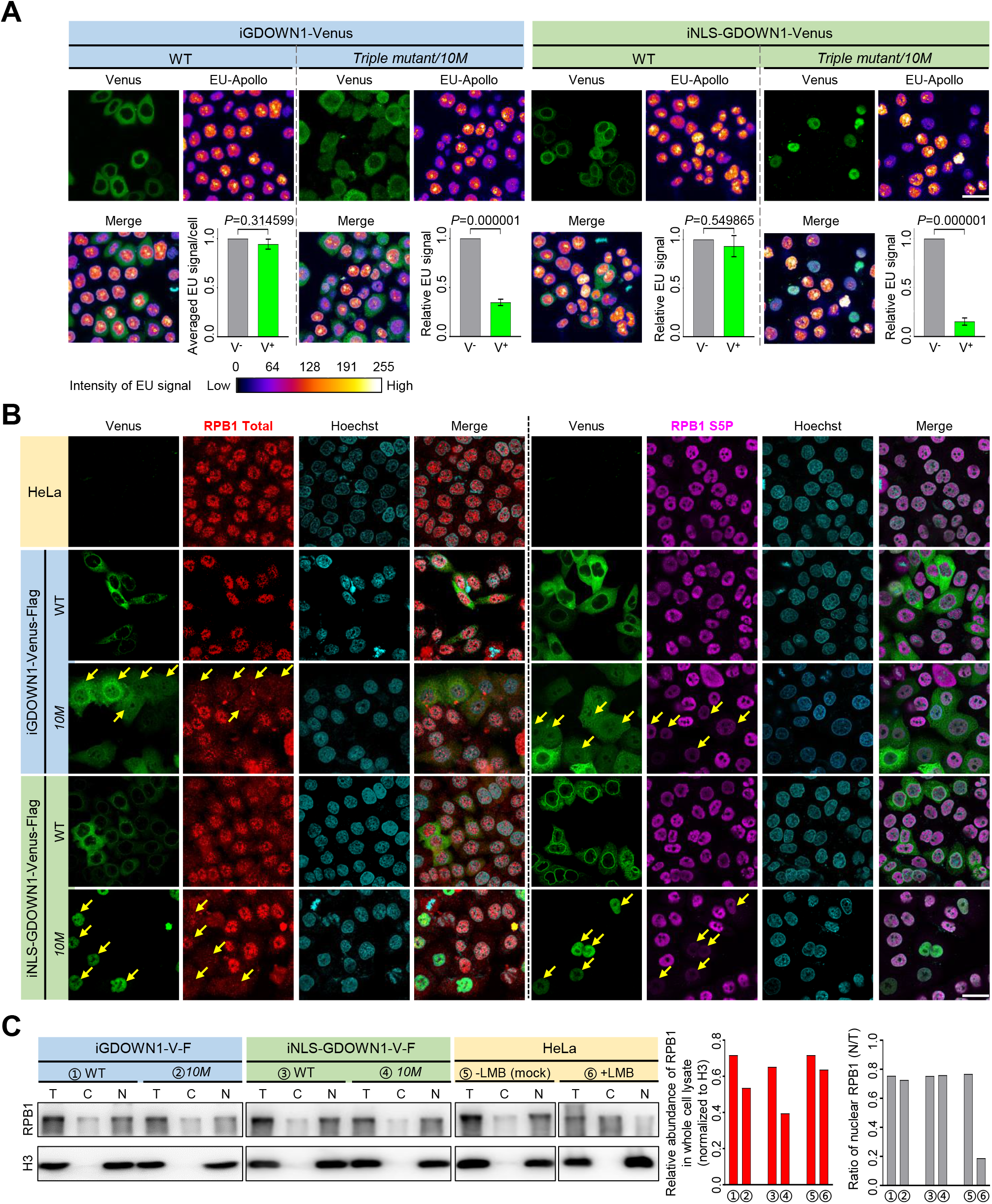
Nuclear GDOWN1 represses global transcription. All the experiments shown in this figure were carried out after four days of doxycycline induction. **A.** Massive accumulation of GDOWN1 in the nucleus caused global transcription repression detected by EU labeling assay. Confocal images were acquired and the EU-Apollo signals were color-coded by imageJ as indicated by the calibration bar shown at the bottom, based on the obtained signal intensity (the original images are shown in Figure S6A). The averaged EU signal/cell value for the Venus^+^ cells (green, V^+^) or for the Venus^-^ cells (gray, V^-^) was shown in the graph at the lower right corner for each indicated cell line. **B.** Nuclear GDOWN1 reduces the levels of total and transcription engaged Pol II. IF experiments were carried out to detect RPB1 levels (total or CTDS5P) in the indicated cell lines. Confocal images were acquired and some representative Venus^+^ cells were pointed out by yellow arrows. **C.** Western blotting analyses of RPB1 in GDOWN1 expressing cells. Each indicated cell line was fractionated to separate cytosol from the nuclei upon harvest, and the cytoplasmic fraction (C), the nuclear fraction (N), and the whole cell lysate (T, total) were further analyzed. Histone H3 served as a nuclear protein control. The RPB1 level in the whole cell lysate relative to that of H3 and the ratio of nuclear RPB1 (N/T) were calculated and shown on the right. The LMB treatment was done as previously described. Nuclear DNA was stained by Hoechst 33342 and all the scale bars represent 30 μm.

### GDOWN1 trans-localizes into the nucleus in response to certain stresses and helps strengthen cellular adaptability

Next, we tested various types of reagents to search for potential cellular stimuli capable of triggering the nuclear translocation of endogenous GDOWN1. No obvious change of GDOWN1’s subcellular localization was observed when cells were treated with the transcriptional inhibitors DRB or Madrasin, the translational inhibitor CHX, or the inhibitors for DNA topoisomerases such as CPT or Doxorubicin (Fig. S7A). Interestingly, we found the treatment of sodium arsenite (NaAsO_2_) reproducibly caused nuclear translocation of GDOWN1. As shown in Figure 7A, NaAsO_2_-induced nuclear translocation of GDOWN1 occurred in a dose-dependent manner and reversed upon NaAsO_2_ removal. Exposure to inorganic arsenite was known to induce global transcription repression (Nelson et al., 2009; Rea et al., 2003) and eventually cause severe cellular toxicity, such as growth inhibition, DNA damage, reactive oxygen species (ROS) production, apoptosis, and autophagy (Tam et al., 2020) and. When cells were treated with 0.5 mM NaAsO_2_ for 30 min (the prevalently used condition in the literature), nearly all cells generated stress granules (SGs) no matter GDOWN1 was competent or knocked out, indicated by the IF signals of G3BP1, a typical SG marker (Fig. S7B). However, under a milder condition (0.1 mM of NaAsO_2_, for 6 hours), which triggered SG formation in a very small fraction of the control cells and in the cells ectopically expressing exogenous GDOWN1-Venus, significantly more of the *GDOWN1 KO* cells already generated SGs (Fig. 7B). EU staining results indicated that this milder NaAsO_2_ treatment strongly downregulated total transcription (Fig. 7C). Furthermore, the viability of *GDOWN1 KO* cells was significantly less than the GDOWN1 competent counterparts (Fig. 7D), indicating that loss of GDOWN1 made the cells hypersensitive to the cell toxicity induced by the low dose of NaAsO_2_ treatment. Taken together, our data demonstrate that the nucleocytoplasmic localization of the native GDOWN1 is switchable in response to NaAsO_2_-induced cellular stress and potentially to other types of unidentified cellular stimuli, and strongly suggest that GDOWN1-mediated transcriptional control contribute to the cellular sensitivity and adaptation to certain stresses.

**Figure 7.**
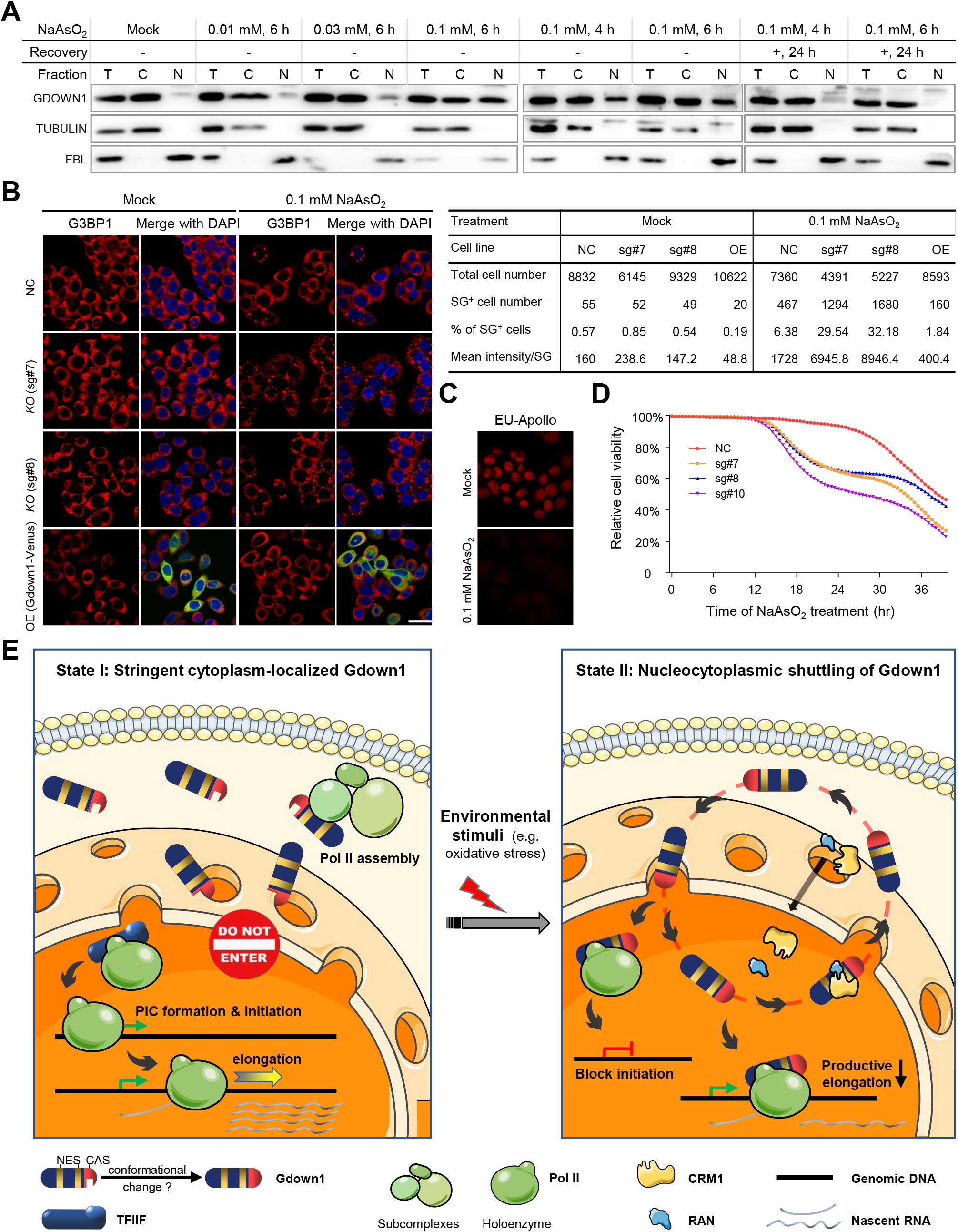
The expression levels of GDOWN1 correlate to the cellular sensitivity to NaAsO_2_ treatment. **A.** Upon NaAsO_2_ treatment, a portion of cellular GDOWN1 was subjected to a reversible translocation into the nucleus. HeLa cells were mock treated or treated with NaAsO_2_ as indicted. In some samples, the cell culture medium was refreshed after treatment to remove NaAsO_2_, and the cells were further cultured for another 24 hours before harvest. Cells were fractionated to separate cytosol from nuclei and the cytoplasmic fraction (C), the nuclear fraction (N) and the whole cell lysate (T, total) were further detected by Western blotting. α-TUBULIN and FBL (a nucleolus protein) were used as markers of the cytoplasmic and nuclear fractions, respectively. **B.** GDOWN1 affected the formation of SGs after NaAsO_2_ treatment. HeLa cells with *GDOWN1* KO (sg#7, sg#8) or the negative control (sg#NC), and cells stably and inducibly expressing iGDOWN1-Venus-Flag (OE) were employed. Each indicated cell line was subjected with NaAsO_2_ treatment at 0.1 mM for 6 hours, and the SGs were detected by immunofluorescence assays using an antibody against G3BP1. Nuclear DNA was stained by Hoechst 33342 and all the scale bars represent 30 μm. Left: representative confocal images; Right: parameters of SGs measured and calculated by Gene 5, based on the images acquired by Cytation 5. **C.** Total transcription level in HeLa cells was repressed upon NaAsO_2_ treatment. HeLa cells treated with 0.1 mM NaAsO_2_ or mock treated were used in EU-Apollo labeling assay. **D.** Loss of GDOWN1 made cells more sensitive to NaAsO_2_ stimulation. Relative cell viability of the indicated cell lines in the presence of 0.1 mM NaAsO_2_ were monitored and calculated by Cytation 5 and Gene 5. **E.** A model summarizing the working and regulatory mechanisms in GDOWN1 (described in the main text).

## DISCUSSION

The appropriate subcellular localization of a protein determines its potential accessibility for certain cellular processes therefore serves as the fundamental premise for executing functions. This study is mainly focused on exploration of Gdown1’s subcellular localization and the associated functional and regulatory mechanisms in mammalian somatic cells. Our results confirmed the cytoplasmic localization of Gdown1 in the cultured cell lines. To demonstrate the nucleocytoplasmic shuttling properties of GDOWN1, we treated HeLa and other types of cells with a specific inhibitor of the nuclear exportin protein CRM1, LMB, with the expectation to observe its nuclear accumulation upon the treatment. Strikingly, it turned out that for all the cell lines tested, GDOWN1 remained its cytoplasmic localization in the presence of LMB, confirmed by both biochemical and cell-based assays. Furthermore, the artificial addition of NLS motifs to GDOWN1 did not efficiently promote its nuclear translocation either. Thus, we conclude that under conventional cell culture conditions, GDOWN1 is strictly locked in the cytoplasm rather than dynamically shuttling between the cytoplasm and the nucleus (Figure 7D), which makes Gdown1 remarkably different from the typical nucleocytoplasmic shuttling proteins.

Our systematic dissection of the intrinsic localization regulatory element(s) in GDOWN1 via mutant analyses let us identify a binary localization regulatory system composed of the functionally coupled NES and CAS motifs. This delicate orchestration between CAS and NES controls the nucleocytoplasmic distribution of Gdown1, guaranteeing the appropriate input of Gdown1 in transcriptional regulation. The facts that both NES and CAS motifs are conservative and the CAS activity seems to be gradually strengthened from lower to higher animals further highlight the essential role of this whole regulatory apparatus/mechanism in controlling Gdown1’s subcellular localization and functions.

In terms of the working mechanisms of the CAS motif, at least it is partially attributed to its participation of anchoring GDOWN1 to the cytoplasmic filament subcomplex of the NPC. NPCs are composed of ∼32 conserved nucleoporin proteins. Besides their central role as nucleocytoplasmic conduits, recent studies have revealed that Nups play an important role in the maintenance of cellular homeostasis through their participation in many cellular activities such as chromatin organization, transcription regulation, DNA damage repair, genome stabilization, and cell cycle control etc. (Raices and D’Angelo, 2021). Therefore, our results support the potential involvement of NPCs in recruitment of GDOWN1 to the nuclear periphery and the resultant cytoplasmic retention, suggesting that the nuclear periphery might be the main workplace for GDOWN1 to execute its cytoplasmic functions. When CAS is fully functional, it locks GDOWN1 in the cytoplasm sufficiently so that the function of NES becomes a backup, which explains the phenomenon that GDOWN1 is insensitive to LMB treatment under this circumstance. Thus, our data suggests that removing or at least alleviating the constraint of CAS would be a prerequisite for licensing GDOWN1’s nuclear translocation and the following transcription regulatory activities. Besides the NPC-anchoring activity, other working mechanisms of the CAS-directed cytoplasmic retention remains to be explored. In addition, the controlling mechanisms for switching off the CAS activity remain unclear. Based on our findings, one reasonable hypothesis is that post translational modifications of the core arginines within CAS or possibly other amino acids nearby might facilitate this switch via causing a conformational change or affecting the interactions of GDOWN1 to its regulatory factors (illustrated in Figure 7D), which is similar to the reported cases in the literature (Ashida et al., 2022; Navarro-Lerida et al., 2021).

Our data demonstrate that mutation of the CAS motif immediately switches GDOWN1 into an LMB-sensitive nucleocytoplasmic shuttling protein and its nuclear abundance is determined by the dynamic balance between its functionally-associated binding partners (such as Pol II) and the CRM1/RAN-mediated nuclear export machinery. This partial translocation of GDOWN1 leads to tremendous changes inside of the nucleus, including the reduction of Pol II and the global transcriptional decrease. The less Pol II, the less active transcription there is, and vice versa, and this mutual feedback causes the drastic decline of cellular transcription level. It was suggested that GDOWN1 was involved in Pol II assembly as well (Ball et al., 2022), therefore its nuclear translocation may also lead to the reduced efficiency of Pol II assembly so that further strengthen its transcription inhibitory effects. Our EU staining results demonstrate that the global transcription is drastically affected, for example, the very strong EU labeled rRNA signals in the nucleoli are dramatically decreased (Fig. 6A). Thus, GDOWN1 also interferes the activity of Pol I and Pol III, while the mechanisms behind this layer of regulation remain unknown.

Recently it was reported that GDOWN1 played a role in facilitating global transcriptional shut down during mitosis and the genetic ablation of *GDOWN1* exhibited mitotic defects (Ball et al., 2022), which is consistent with GDOWN1’s stringent localization in the cytoplasm during the interphase. Our discovery of GDOWN1’s nuclear translocation upon cellular stresses further expands the context in which GDOWN1 plays an essential role in global transcription repression. The cells without GDOWN1 are much more sensitive to cellular stresses, emphasizing that GDOWN1 is a crucial factor in maintaining cellular homeostasis, and further studies are needed to explore GDOWN1’s functions in the cytoplasm and to identify more cellular situations that trigger its nuclear translocation. Taken together, this work uncovered GDOWN1’s new functions and switchable localization in mammalian somatic cells and shed a light on a new connection of the global transcriptional regulation and cellular stress adaptation.

## MATERIALS AND METHODS

### Key resources table

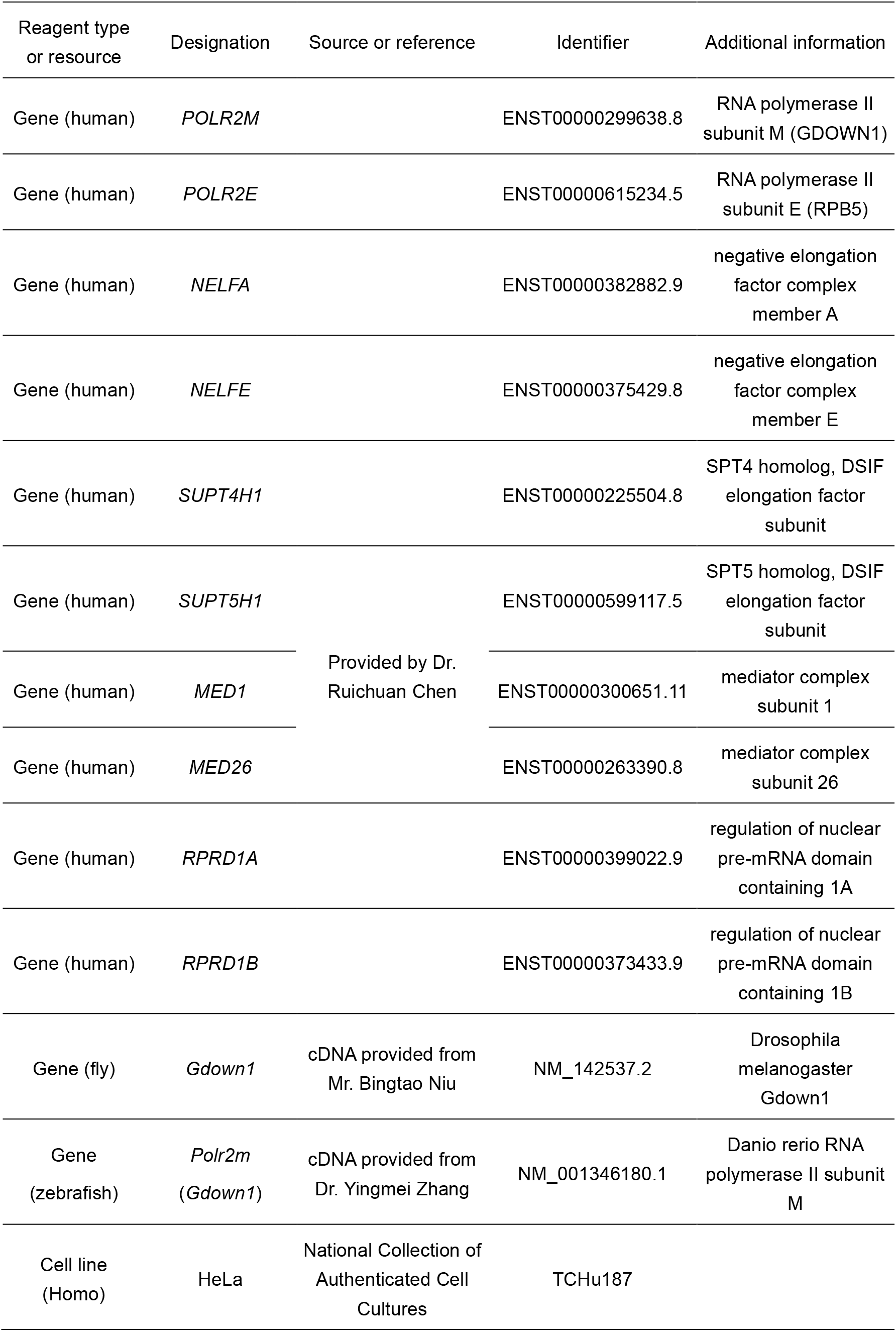
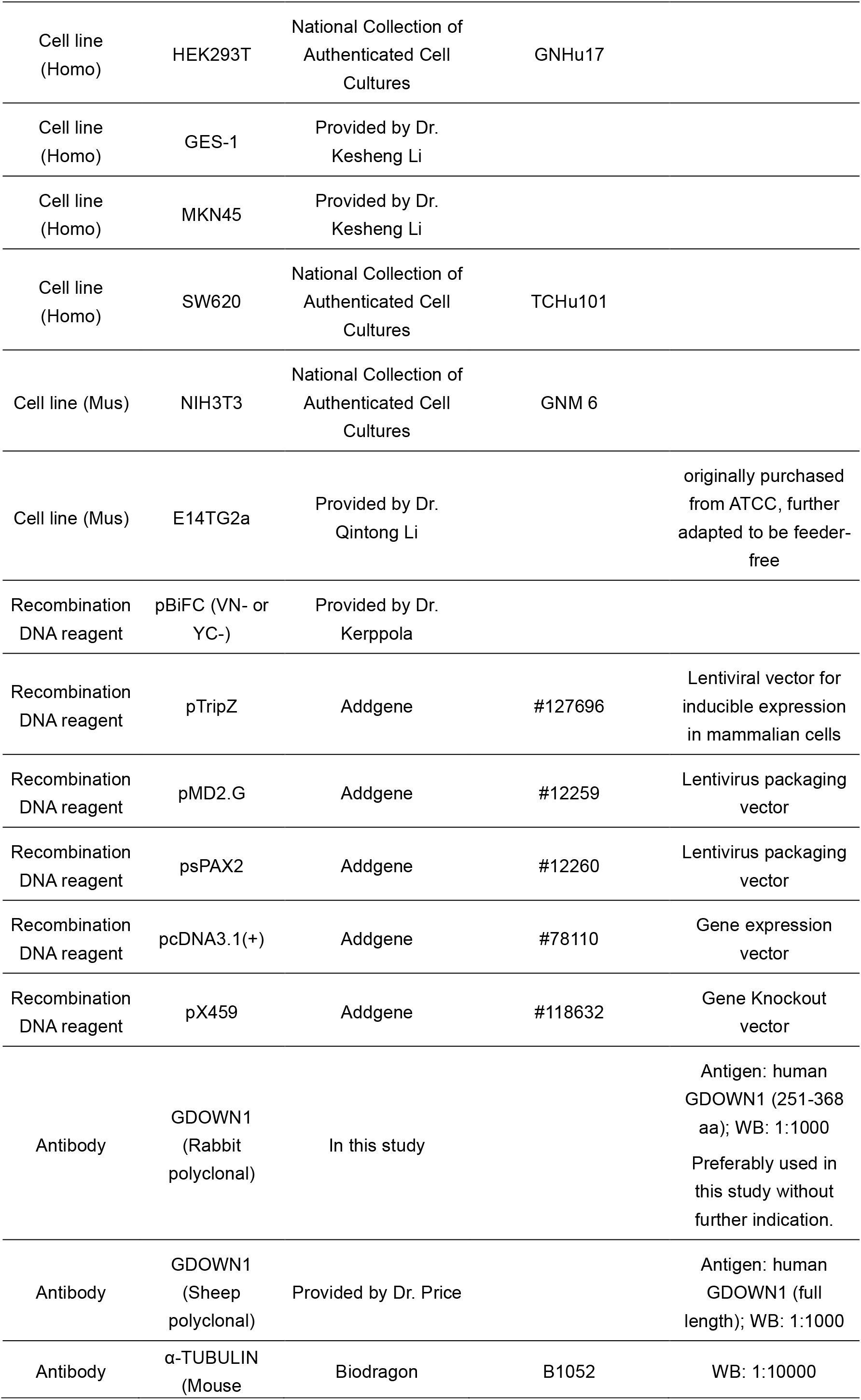
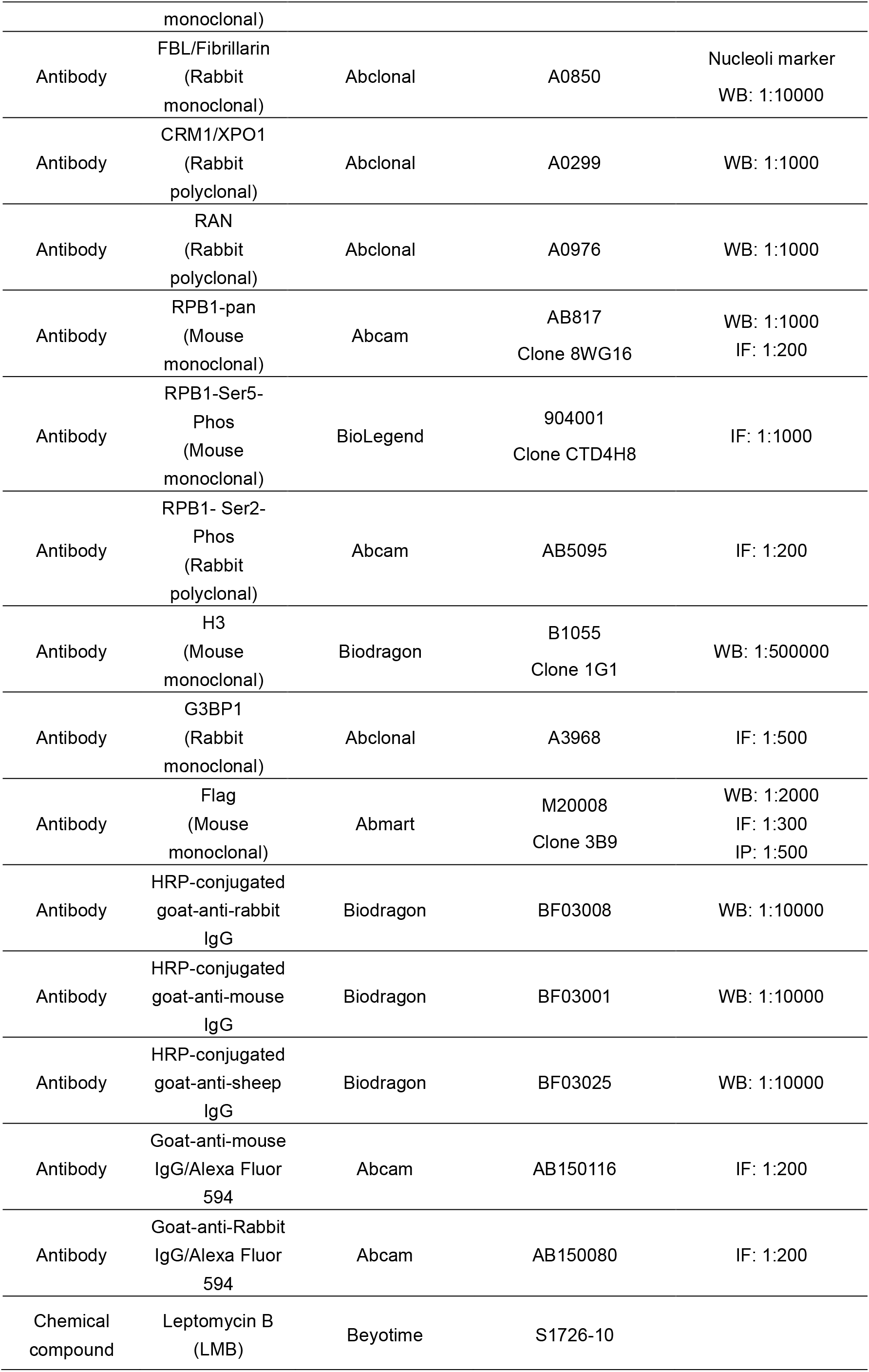
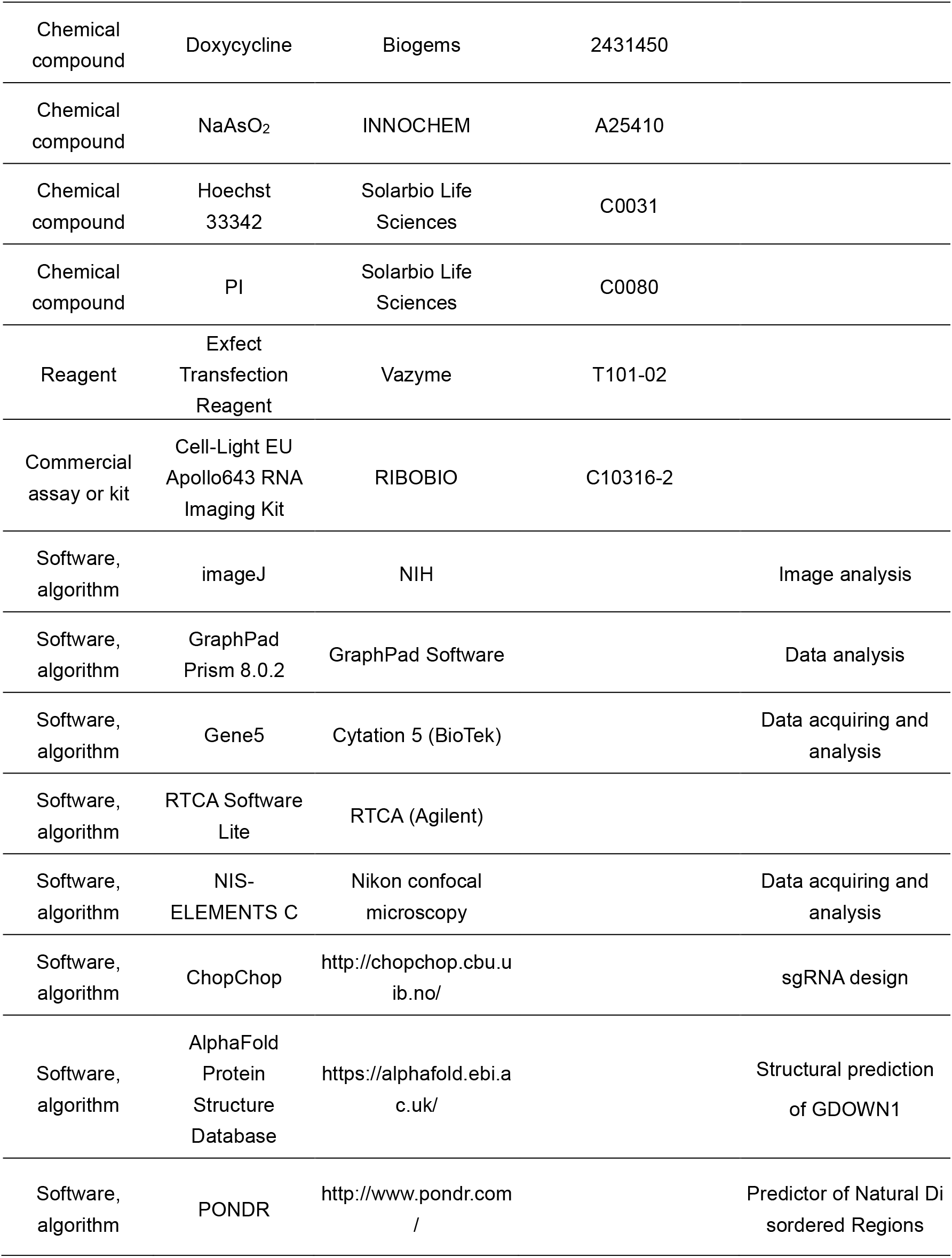

### Cell culture, transfection, and drug treatment

HeLa cells and all the other cell lines except for E14Tg2a were cultured in Dulbecco’s Modified Eagle’s Media (Gibco, 12800-017) supplemented with 10% Newborn Calf Serum (Biological Industries, 04-102-1A) and pen/strep. The mouse embryonic stem cell line,

E14Tg2a, (gift of Dr. Qintong Li in Sichuan University) were cultured in Dulbecco’s Modified Eagle’s Media supplemented with 15% Fetal Bovine Serum (Gemini Bio-products, 900-108), 1x non-essential amino acids (Gibco, 11140-035), 200 mM L-glutamine, 0.1 mM β-mercaptoethanol, 10^3^ U/mL leukemia inhibitory factor (LIF, purified in lab), and pen/strep. All the dishes or coverslips used for culturing E14Tg2a cells were pretreated with 0.5% gelatin. All cells were maintained at 37°C, 90% humidity and 5% CO2. Plasmid transfections were carried out using Exfect Transfection Reagent according to the manufacturer’s protocol. 0.25 μg plasmid was used for transfecting one well of cells in a 24-well cell culture dish and normally confocal microscopy images were taken at 24 hours post transfection.

For samples treated with LMB, 20 nM final concentration of LMB was added to the culture medium at 18 hours post transfection and incubated for 6 hours before data collection (or mock treated with an equal volume of ethanol). For NaA_S_O_2_, DRB, CHX, Madrasin, Tubercidin, CPT and Doxorubicin treatment, the drug was added to the complete medium to the indicated final concentration and incubated with cells for indicated timing. Cells were washes for three times with PBS to remove the drug before further operation was pursued.

### Construction of plasmids and stable cell lines

The pBiFC-Flag-VN (1-172 aa of Venus) or pBiFC-Flag-YC (173-238 of YFP) plasmids (gifts from Dr. Tom Kerppola, University of Michigan) were used as parental vectors for generating all the indicated BiFC plasmids. The coding sequences of human *MED1*, *MED26*, and *SPT5* genes were PCR amplified from plasmids (gifts from Dr. Ruichuan Chen, Xiamen University, (Lu et al., 2016)) and *GDOWN1* (also namely *POLR2M*, NM_015532.5) and other genes were all amplified by RT-PCR used cDNA templates generated from HeLa cells. The *Gdown1* genes in Danio rerio (NM_001346180.1) and Drosophila melanogaster (NM_142537.2) were cloned from cDNA samples generated directly from animal lysates. Total RNA was extracted by MolPure Cell/Tissue Total RNA Kit (YEASEN, 19221ES50) and the cDNA was synthesized using 1^st^ Strand cDNA Synthesis SuperMix (YEASEN, 11141ES60). The purified RT-PCR products were double digested by BamHI and XbaI (NEB) and then ligated into pBiFC-Flag-VN or -YC vectors by T4 DNA ligase or when these two restriction enzymes had cut sites within the cDNA sequences, the PCR products were assembled into pBiFC-Flag-VN or -YC vectors via homologous recombination using ClonExpressII One Step Cloning Kit (Vazyme, C112-02). The two NLS motifs in plasmid namely Flag-NLS-GDOWN1-NLS in Figure 1A were adopted from pX459 originally constructed from Dr. Feng Zhang’s lab in MIT (Ran et al., 2013). The three NLS motifs in VN/YC-3xNLS-GDOWN1 plasmids shown in Figure 1D were cloned from the CDS sequences of human *RYBP* gene corresponding to amino acids 1-94 (Tan et al., 2017). The truncated fragments of human *GDOWN1* were amplified using the full-length CDS as a template and further used to construct pBiFC-based GDOWN1 mutants. Point mutations were introduced by designing long PCR primers containing the designated mutated sequences and then amplified the fragments by regular PCR or bridging PCR as needed. The information of the amino acids for mutagenesis was shown in Figures 2A and 3A. The above pBiFC-based plasmids series were applied in both BiFC assays (directly monitoring BiFC signals) or in immunofluorescence assays (detection via Flag antibody) as indicated in the figure legends. For generating the GDOWN1-Venus plasmid series, pcDNA3.1(+) was used as a parental vector. Full-length, wild type GDOWN1 was amplified from the above pBiFC vector and ligated into pcDNA3.1-Venus plasmid (previously constructed in lab). The NES and/or CAS mutant fragments were PCR amplified from the above pBiFC plasmids expressing the corresponding mutant GDOWN1 and further amplified by bridging PCR and then assembled into the pcDNA3.1-Venus plasmid.

*GDOWN1* KO HeLa cells were generated via CRISPR-Cas9 technology. The sgRNAs were selected according to the information provided by ChopChop. The targeting sequences of sgRNAs are listed in the table down below. The pX459-sg*GDOWN1-*#1/#7/#8/#10 plasmids were constructed and transfected into HeLa cells and the cells were selected with 0.5 μg/mL puromycin starting from 48 hrs post transfection. After 5 days of selection, the survived cells were pooled and further verified by sequencing and WB. For cells transfected with pX459-sg*GDOWN1-*#1, the pooled cells after puromycin selection were replated in a p100 cell culture dish at a density of 2000 cells per dish. After 15 days of culture, single colonies were picked to a 96-well plate and further expanded. Genomic DNA was isolated and PCR amplified using verification primers shown in the table. The PCR products were gel purified and sent for sequencing (Tsingke Biotechnology).

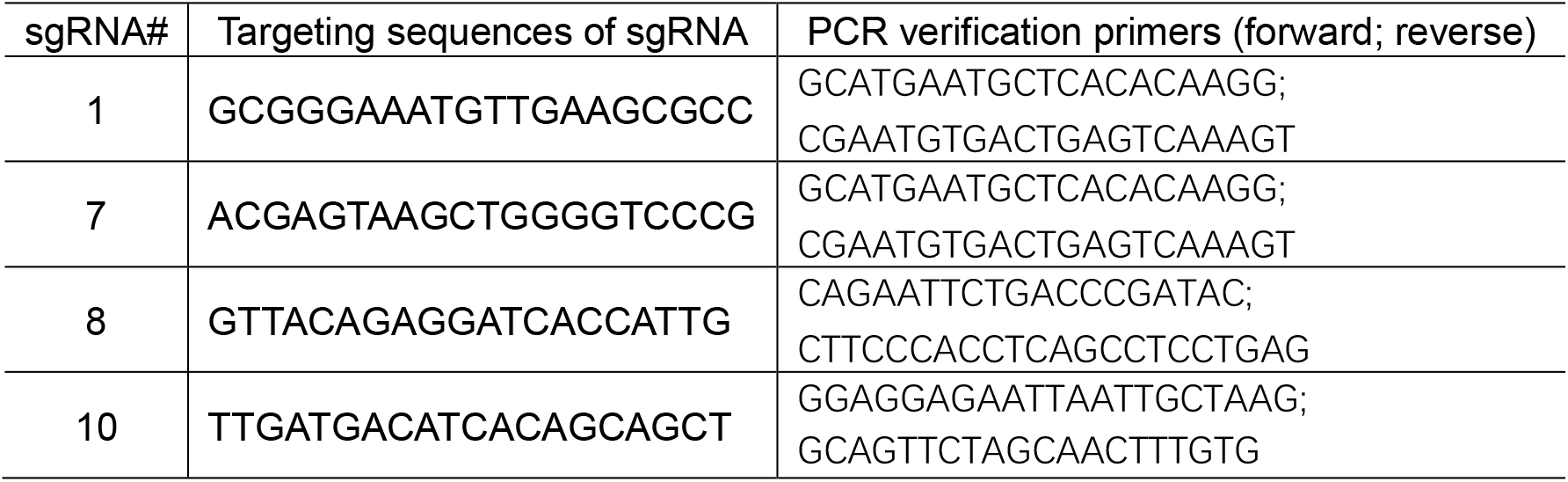

For generating HeLa cell stably expressing GDOWN1, the lentiviral expression vector, pTripZ was used as a parental vector and the fragment, Venus-Flag, was initially inserted into pTripZ empty vector to replace the original shRNA expression cassette. The wild type or mutant GDOWN1 fragments were PCR amplified from the constructed pcDNA3.1-based plasmids (for the ones with NLS addition, the sequences of SV40-NLS were attached to the N-terminus of the corresponding primers) and were further inserted in between the TRE-CMV promoter and the Venus gene to obtain iGDOWN1-Venus-Flag plasmids (“i” stands for inducible). For viral packaging, HEK293T cells cultured in a 6 well plate were transfected with 1 μg pMD2G, 2 μg pAX2 and 3 μg pTripZ-GDOWN1-Venus (WT or mutant) and medium was refreshed at 6 hrs post transfection. The viral stock was harvested after 72 hrs and further infected HeLa cells for 12 hrs. Cells were recovered for 1 day and further subjected for puromycin selection (0.5 μg/mL) for 14 days. The survived cells were pooled and the inducible expression of the GDOWN1-Venus-Flag proteins was verified by WB with a Flag antibody after adding 2.5 μg/mL doxycycline for 12 hrs.

### BiFC assays

For BiFC assays, HeLa cells were grown on coverslips in 24-well cell culture dishes and 0.25 μg of each pBiFC plasmid (VN- or YC-) was used for co-transfection per well. At 24 hrs post transfection, BiFC complexes in transiently expressing cells were fixed with 4% formaldehyde for 20 min at room temperature, washed with PBS, stained with 1 μg/mL of Hoechst 33342 for 10 min, washed with PBS, and visualized in PBS.

### Immunofluorescence and data analyses

For immunofluorescence assays, cells were grown on coverslips, fixed with 4% formaldehyde for 20 min at room temperature, washed three times with PBS, dehydrated with 90% methanol at -20°C for 30 min, permeabilized with 0.5% Triton X-100 at room temperature, washed three times with PBS, incubated with 5% BSA for 1 hr at room temperature. Cells were then incubated with Flag antibody (1:200 diluted in TBST) for 12 hrs at 4°C. After being washed for three times with TBST, the cells were subjected for secondary antibody incubation for 1 hr at room temperature. The cells were further stained with 1 μg/mL of Hoechst 33342 for 10 min, washed with PBS, and visualized in PBS.

For stress granules statistics, cells were grown on coverslips in 24-well cell culture dishes (inducible cell lines were pre-induced for 1 days) and Mock or 0.1 mM NaAsO_2_ were used to treat cells. At 6 hrs post treatment, cells were washed with PBS 3 times, and used G3BP1 antibody to do IF as above description. Confocal images were taken and the images were acquired using NIS-ELEMENTS C software. For data quantification, cells were monitored in a Cytation 5 live cell detection system using a 10×objective. At least 4 ROIs were randomly selected from each well, and all of images were acquired using Gene 5 software using the same parameters, and combined for further data analyses. The count of total cells (Hoechst 33342 signals were used as an indicator), spot number in every cell and the mean fluorescence intensity of each spot (spots of G3BP1 signals were used as an indicator) in each ROI was calculated using the built-in tools (Automatic cell count, spots count and subpopulation analysis) in Gene 5 software. the cells contained more than 1 spot were counted as an SG^+^ cell.

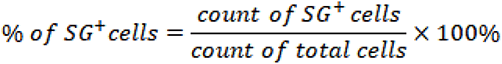

### Co-immunoprecipitation

Cells stably expressed GDOWN1-Venus-Flag in p100 dish were lysed by adding 500 μL of lysis buffer [20 mM Tris-HCl, pH 8.0, 150 mM NaCl, 0.5 mM ethylenediaminetetraacetic acid, 1% NP40, 1% Triton X-100, 50 mM NaF, 1 mM Na_3_VO_4_ (activated), 0.1 mM PMSF, Protease Inhibitor Cocktail (Bimake, B14012)] and incubated for 30 min at 4°C with rotation. The lysate was incubated with 25 μL of anti-Flag magnetic beads (Bimake, B26101) for 12 hrs at 4°C on a rotator. The beads were washed five times with lysis buffer, and then resuspended with 5X loading dye (250 mM Tris-HCl, pH 6.8, 10% SDS, 50% glycerinum, 5% β-mercaptoethanol, 0.1% bromophenol blue). The samples were boiled at 100°C for 10 min and used for SDS-PAGE analyses.

### Confocal Microscopy

Confocal images were obtained using a 100x oil objective (N.A. 1.45) on a Nikon A1R+ Ti2-E laser scanning microscope equipped with a GaAsP Multi Detector Unit. Images were acquired from at least 4 randomly selected fields using NIS-ELEMENTS C software. For data quantification of GDOWN1’s nucleocytoplasmic distribution, images of at least 4 fields from each treatment were randomly selected and imageJ was employed to acquire fluorescence intensity of Venus signals in the entire cells (total signals) and in all the nuclei (nuclear signals) from all the transfected cells (Hoechst 33342 signals were used as an indicator to define the nuclei) and the cytoplasmic signals were calculated by subtracting the nuclear signals from the total signals. The proportion of the cytoplasmic signals (green) and the nuclear signals (blue) were calculated and plotted. For statistics analyses, the calculated values from imageJ were further processed to obtain the P values via t-test using the built-in tools in Graphpad Prism8.

### EU-Apollo assay

For EU-Apollo assays, parental HeLa cells or derived HeLa stable cell lines were grown on coverslips in 48-well cell culture dishes and 0.25 μg/mL doxycycline was used for induction as indicated in figure legend. 250 μM of EU was added to the culture medium at 20 min before cell harvest, then the cells were washed with PBS for 3 times, then fixed with 4% formaldehyde for 20 min at room temperature and followed by quenching with 1 mg/mL glycine solution for 1 min. Then the cells were permeabilized with 0.5% Triton X-100 at room temperature, washed twice with PBS, followed by incubation with 0.5x Apollo 643 staining solution in the Cell-Light EU Apollo 643 RNA Imaging Kit at room temperature for 10 min. After being washed with 0.5% Triton X-100 for 3 times, the cells were further stained with 1 μg/mL of Hoechst 33342 for 10 min, washed with PBS, and finally visualized in PBS. Confocal images were acquired as previously described. For data quantification, images of at least four fields from each treatment were randomly selected and each cell was separated into either Venus positive or Venus negative group based on the fluorescence intensity of Venus. The fluorescence intensity of EU-Apollo signal was measured cell by cell with imageJ and the averaged EU-Apollo signal for each group was calculated and plotted in bar graphs. For statistical analyses, the calculated averaged EU-Apollo values in each field were further processed to obtain the P values via t-test using the built-in tools in Graphpad Prism8.

### Cell fractionation and the quantitative analysis of Western Blot

Freshly harvested cell pellet was resuspended with five volumes of cytoplasmic extraction buffer (20 mM Hepes, 1 mM ethylenediaminetetraacetic acid, 10 mM KCl, 2 mM MgCl_2_, 0.1% Nonidet P-40, 1 mM DTT, 0.1 mM PMSF, Protease Inhibitor Cocktail), and incubated at 4°C for 30 min. The completion of this step was monitored and confirmed by morphology checking under microscope. The cell lysate was centrifuged at 1500 rpm for 3 min at 4°C and the supernatant was saved as the cytoplasmic fraction. The remained cell pellet was washed for three times and further resuspended with cytoplasmic extraction buffer. These resuspended nuclei samples were used as the nuclear fraction (containing both the soluble nucleoplasm and insoluble chromatin). 5x loading dye was added into the above cytoplasmic (C) and nuclear (N) fractions to generate 1x samples for SDS-PAGE and WB analyses.

For data quantification, imageJ was employed to acquire the IntDen value (integral optical density) of each band in the obtained WB images.

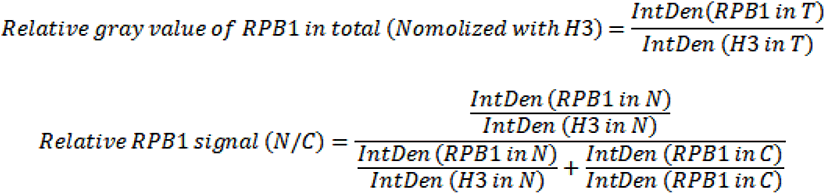

### Live cell analyses and data analyses

For live cell analyses via Cytation 5 (BioTek), cells were plated in a 48-well cell culture dish 24 hrs before the treatment. For the results shown in Figure 7D, cells were incubated with the complete medium supplemented with 0.2 mM NaAsO_2_, 0.1 μg/mL Hoechst33342 and 1 μg/mL PI, then immediately analyzed using a live cell analyzer. Four ROIs from each well were randomly selected and images were acquired using Gene 5 software using the same parameters. The images were stitched together for further data analysis. The count of total cells (using the Hoechst 33342 signal as an indicator) or dead cells (using PI signal as an indicator) in each ROI was calculated using the built-in tools (Automatic cell count and subpopulation analysis) in Gene 5 software. The calculated values were further processed using Graphpad Prism8.

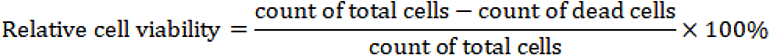

For the results shown in Figure 5B-ii, cells were grown in 6-well cell culture dishes and induced by 0.25 μg/mL doxcycline for 0, 3, 6, 9 days, respectively. At times for harvest, cells were fixed with 4% formaldehyde for 20 min at room temperature, washed with PBS, and visualized in PBS. Images of at least 4 randomly selected ROIs from each well were acquired using Gene 5 software. For data analysis, the count of total cells (using the Hoechst 33342 signal as an indicator) and the Venus positive cells, and the fluorescence intensity of the Venus positive cells was calculated using the built-in tools (Automatic cell count and subpopulation analysis) in Gene 5. The obtained data were further processed in Graphpad Prism8 to export figures.

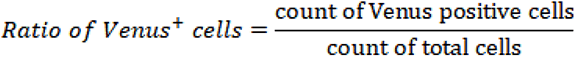

For live cell analyses using RTCA in Figure 5B-iv, cells were seeded in a special 16-well plate supplied by Agilent, and cultured in the equipment set inside of a cell culture incubator. The cell index and the slope of which were acquired by RTCA Software Lite along the growth of cells. The obtained values were further processed in Graphpad Prism8 for data export.

### Cell counting and Flow cytometry analyses (FACS)

For results in Figure 5B-iii, equal amounts of various cells were plated on day 0 and cultured with complete medium. Two experiments were performed at the same time, with the cells seeded in 12-well plates for cell counting and cells in 6 well plates for FACS. Cells were incubated with 0.25 μg/mL doxcycline for 0, 3, 6, 9 days, respectively. When the confluency reached 90%, one third of the cells were passage into a new cell culture dish. The number of cells on day 3, 6 and 9 was counted using a cell counter and the average value was obtained based on three independent readings. The finalized cell counts shown in figure 5B-iii were calculated based on formula below. Cell count=the averaged cell counter reading x 3^the number of passages^.

For FACS analyses, the cells were harvested on day 0, 3 and 9 after doxycycline induction. To gather the dead cells, the culture medium was centrifuged at 1000 x g for 5 min and collect the cells at the bottom of the tubes. Then the adherent cells in the plates were trypsinized (0.25% trypsin) and collected via centrifugation at 800 x g for 5 min. Both the attached cells and the dead cells recovered from the medium in the same sample were combined and resuspend with 500 μL PBS, and further incubated with DAPI (final concentration of 5 μg/mL) at dark for 15 min. Then the cells were immediately handled in the Flow Cytometer (LSRFortessa^TM^, BD) for detection. For each sample, a minimum of 10,000 cells were analyzed with FlowJo 7.6 software. To establish appropriate gating parameters for accurately distinguish DAPI positive dead cells from live cells, we generated a sample by mixing 2/3 of live cells with 1/3 of formaldehyde fixed dead cells, and the scope was delimited after DAPI staining under the same experimental conditions.

## Acknowledgements

The authors acknowledge Dr. David Price (the University of Iowa) for providing Gdown1 antibody made in sheep (Cheng et al., 2012) and for the helpful comments and discussion from his group; Dr. Ruichuan Chen (Xiamen University), Dr. Qintong Li (Sichuan University), Dr. Yingmei Zhang and Bingtao Niu (Lanzhou Univeristy) for providing necessary plasmids, cells and animals; many members of Cheng laboratory for their helpful discussion; Yaojia Wang for her help of revising our model diagram. We also thank the core facilities in the School of Life Sciences, Lanzhou University for providing high-quality instruments and service for our confocal microscopy experiments. We apologize to our colleagues in transcription field whose work was not discussed adequately owing to space constraints.

## Author Approvals

All authors have seen and approved the manuscript, and that it hasn’t been accepted or published elsewhere.

