## Supplemental figures for "Lifting the ban on nuclear import activates Gdown1-mediated modulation of global transcription and facilitates adaptation to cellular stresses"

**SUPPLEMENTARY FIGURES AND LEGENDS**

**Figure S1. A.** Verification of Gdown1 antibodies used in this study. Human *GDOWN1* locus and the targeting positions of sg#1, #7, #8 and #10 are shown as yellow bars. HeLa cells were transfected with pX459- sg#1, sg#7, sg#8 or sg#10 plasmid, respectively, and subjected to puromycin selection. The survived cells were subjected to genomic DNA isolation, PCR amplification and sanger sequencing. Part of the sequencing results surrounding the cleavage sites in the indicated samples are shown as screenshots of the chromas images, with PAM sites marked in green boxes and the edited sequences at the cleavage sites marked in red boxes. Clone #F7 was a single colony selected from pX459-sg#1 transfection and the rest samples were cell pools collected after pX459- sg#7, sg#8 or sg#10 transfection and selection. All the samples were also verified via WB using two Gdown1 antibodies generated from rabbit or sheep, respectively, shown on the right. **B.** The indicated cell line was fractionated to separate cytosol from nuclei and the cytoplasmic fraction (C), the nuclear fraction (N) and the whole cell lysate (T, total) were further detected by WB as described in Figure 1B. **C.** HeLa cells were subjected to either mock or LMB treatment (as described in Figure 1) before further fractionation. WB analyses were carried out to check the subcellular localization of GDOWN1 using a Gdown1 antibody

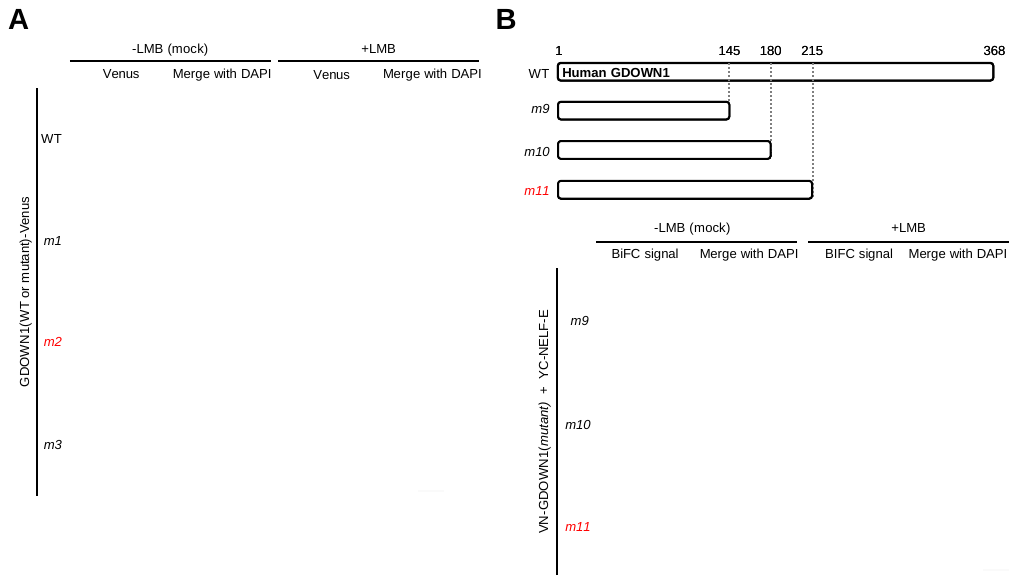

**Figure S2.** **A.** Detection of the subcellular localization of GDOWN1 (wild type or mutants)-Venus. HeLa cells were transiently transfected with indicated plasmids, induced by 2.5 μg/mL doxycycline for 12 hours, further subjected to either mock or LMB treatment, and detected by a confocal microscope. **B.** Identification of the NES1 motif in the BiFC reporter system as described in Figure 2A-B. The details for the truncation mutants used are shown on the top. Nuclear DNA was stained by Hoechst 33342 and all the scale bars represent 30 μm.

**Figure S3. A.** Detection of the subcellular localization of GDOWN1 (wild type or its mutants)-Venus by IF. The details of the indicated truncation mutants are shown on Figure 3A and the cell treatments were performed as described in Figure S2A. After cell harvest, IF experiments were carried out using a FLAG antibody. **B.** The details of the truncation mutants used in Figure S3C. **C.** Identification of NES2 and CAS motifs in the BiFC reporter system as described in Figure 3A-B. Nuclear DNA was stained by Hoechst 33342 and all the scale bars represent 30 μm.

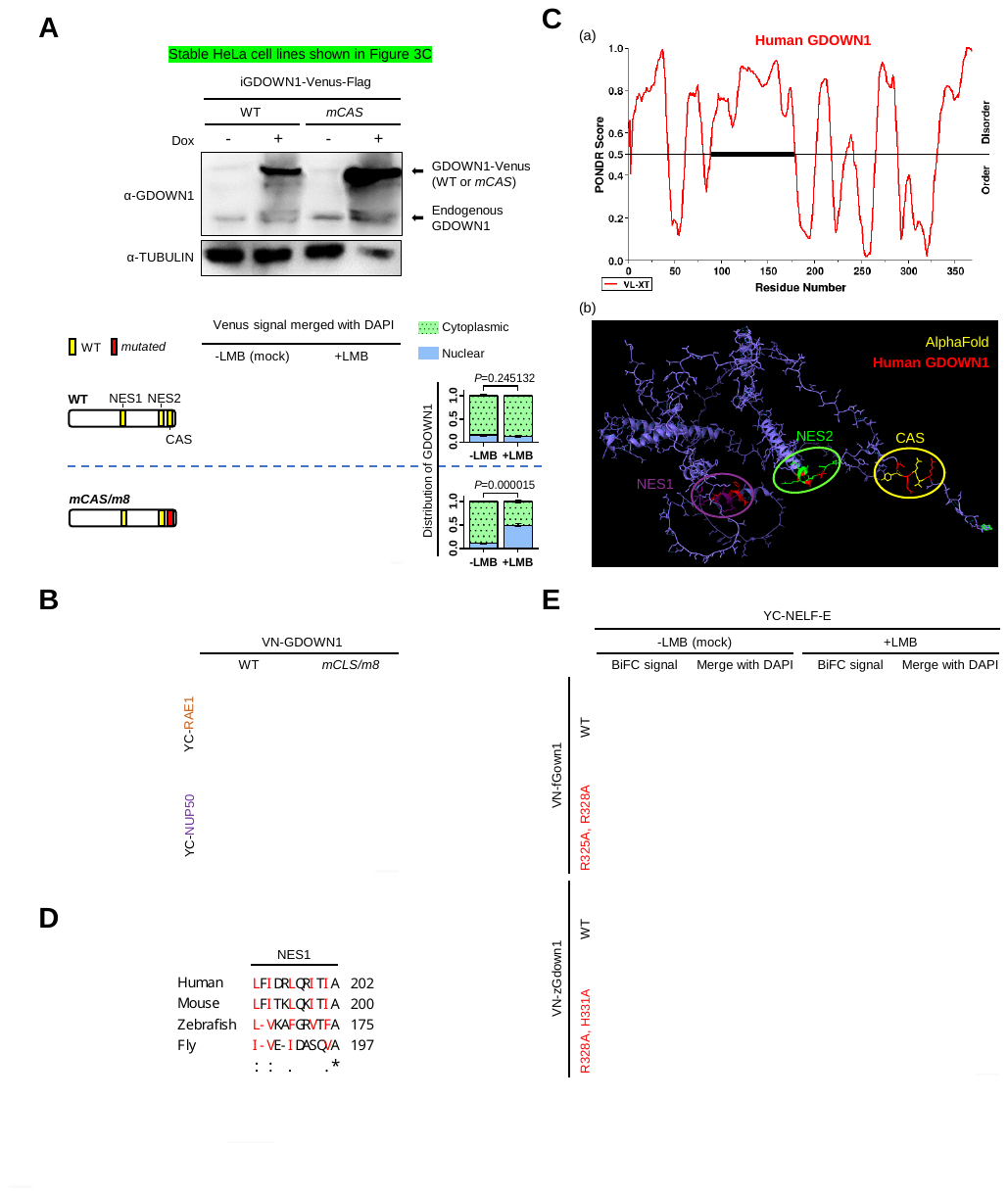

**Figure S4. A.** Top panel: HeLa cells stably transfected the wild type GDOWN1-Venus or the CAS mutant fused with Venus (in pTripZ backbone) were verified by WB using Gdown1 antibody. Bottom panel: The cells were treated with 2.5 μg/mL doxycycline for 12 hours to induce the expression of GDOWN1-Venus, and then subjected to mock or LMB treatment as described in Figure 1. Data collection, processing and statistics analyses were carried out the same as described in Figure 4A. **B.** The enrichment of GDOWN1 at the nuclear pore region was regulated by the CAS motif. Detect the interaction between GDOWN1 or its CAS mutant and NPC components as described in Figure 3D, except that the fluorescence intensity of the BiFC signals were adjusted to similar levels for better observation. **C.** Structural prediction of human GDOWN1. (a) distribution of the ordered and disordered regions within human GDOWN1 using PONDR (http://www.pondr.com/). (b) the structure of human GDOWN1 predicted by AlphaFold2 (https://alphafold.ebi.ac.uk/). NES1, NES2 and CAS were marked with purple, green and yellow circles, respectively, and the mutated amino acids in this study were marked with red. **D.** Sequence alignment of the putative NES1 regions of Gdown1 proteins from the indicated species. “*”—identical in all species analyzed; “:”—highly conserved; “.”— moderately conserved. **E.** BiFC assays monitoring the interaction signals between NELF-E and fly (f) or zebrafish (z) Gdown1, wild type or CAS mutant as indicated. Nuclear DNA was stained by Hoechst 33342 and all the scale bars represent 30 μm.

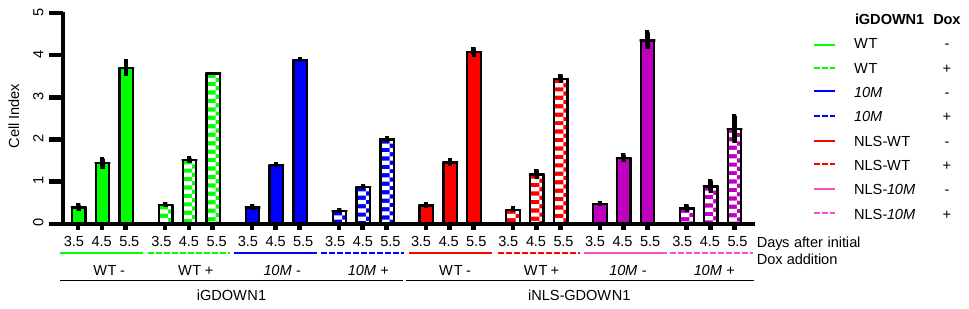

**Figure S5.** Cell index at the indicated time points in Figure 5B-(iv) were further demonstrated in bar graph via Graphpad Prism8.

**Figure S6. A.** The original confocal images from EU-labeling experiments shown in Figure 6A (top). The bottom part shows the confocal images about the IF results for the indicated cell lines detecting Ser-2 phosphorylated RPB1 (CTDS2P) after 4 days of doxycycline induction. Nuclear DNA was stained by Hoechst 33342 and all the scale bars represent 30 μm. **B.** Supplementary WB information of Figure 6C. α-TUBULIN and FBL were used as markers of the cytoplasmic and nuclear fractions, respectively.

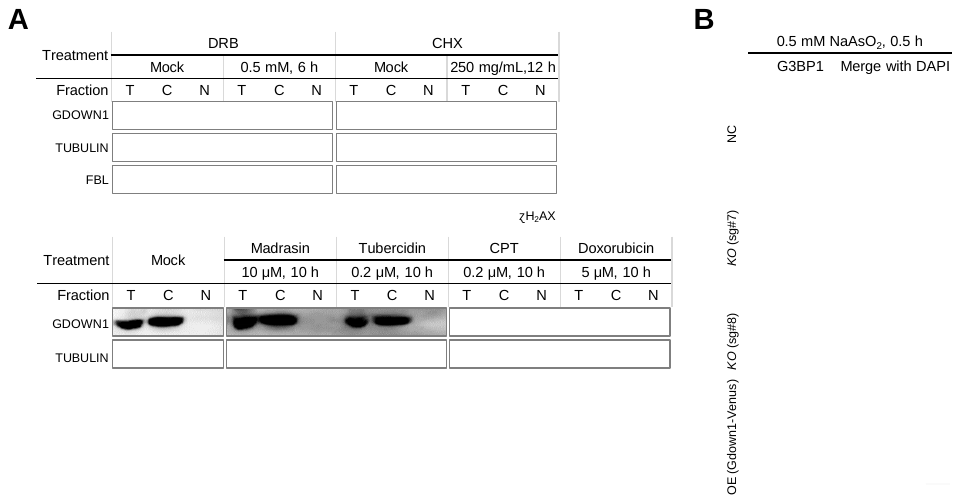

**Figure S7. A.** Subcellular localization of GDOWN1 upon various types of treatments, detected by cell fractionation followed by WB analyses. HeLa cells were mock treated or treated with DRB, CHX, X-ray, Madrasin, Tubercidin, CPT or Doxorubicin as indicated. After treatment, cells were fractionated to separate cytosol from nuclei and the cytoplasmic fraction (C), the nuclear fraction (N) and the whole cell lysate (T, total) were further detected by WB. **B.** Each indicated cell line was treated with 0.5 mM NaAsO_2_ for 30 mins, and the formation of SGs was detected by IF using a G3BP1 antibody. Nuclear DNA was stained by Hoechst 33342 and all the scale bars represent 30 μm.
